# The fodder grass resources for ruminants: A indigenous treasure of local communities of Thal desert Punjab, Pakistan

**DOI:** 10.1101/796227

**Authors:** Humaira Shaheen, Rahmatuallah Qureshi, Mirza Faisal Qaseem, Piero Bruschi

## Abstract

Indigenous people have been using their regional grasses for rearing their animals for centuries. The present study is the first recorded traditional knowledge of grasses and feeding system for livestock from the Thal desert in Pakistan. Snowball method was used to identify key informants. Information was collected from 232 informants from six districts of Thal Desert through semi-structural questionnaire and site visits. The data was analyzed through Smith’s salience index and Composite Salience using ANTHROPAC package in R software. On the whole 61 grasses were recorded from the study area and most of the species belongs to the Poaceae family (52 species). Based on palatability grasses were categorized into three major groups i.e. (A) High priority, (B) Medium priority and (C) Low priority. Species in Group A, abundantly present in the study area, highly palatable forage for all ruminants. 232(141M +91W) local informants were interviewed. Informants were grouped into three major age categories: 20–35 (48 informants), 36–50 (116 informants) and 51–67 years (68 informants). ANTHROPAC frequency analysis conformed the Smith’s salience index and Composite Salience; *Cynodon dactylon* was the favorite species (6.46 SI, 0.6460 CS) followed by *Cymbopogon jwarancusa* (5.133 SI, 0.5133 CS) and *Sorghum* sp. was the third most salient species (5.121 SI, 0.5121 CS). Grasses were mostly available during the season of August and October and had also ethnoveterinary importance. This document about the traditional feeding of livestock from Thal Desert can strengthen the value of conserving our traditional knowledge, which was poorly documented before.

## Introduction

In rural areas of Pakistan, agro-pastoral activities play a crucial role in the development of the local economy, accounting for more than half of the total agricultural income and 10.6% of the national GDP [1]. These activities are particularly important in the economy of the country’s desert regions where land cultivation is difficult and livestock is the main and often unique survival strategy and income source for the local communities. Moreover, milk and meat production may counteract the impact of climatic unpredictability on fluctuations in food availability, especially in areas facing frequent crop shortages. According to data reported by Farooq et al. [2], in Pakistan 8.1% of buffaloes, 13.5% of cattle, 15.3% of sheep and 14.4% of goats are raised in desert districts. However, husbandry in these areas is often an uncertain and low-paid activity; shortage of fodder as a result of severe climatic conditions, high rate of diseases, limited availability of veterinary services and poor access to animal vaccination are important constraints limiting the local livestock productivity [2]. The sustainable production of livestock under harsh climatic conditions needs efficient strategies for improving fodder utilization and management [3]. From this perspective, traditional knowledge can be an important source of information on local wild forage resources and on their nutritive properties. Several studies have shown that smallholder farmers in many parts of the world have a deep practical knowledge about the importance and quality of plants used to feed animals. Ethnobotanical investigations on fodder plants have been carried out in Africa [4–6], Brazil [7], India [8, 9] and China [10–12]. Many studies throughout the world highlight the diverse and abundant use of grasses and sedges as fodder [12, 13] [7, 8]; grasses and sedges are generally reported to be palatable and highly productive resources and to have a high forage potential especially in arid and semiarid areas [7].

Previous studies have shown that Thal is rich in grasses and sedges [14]; most of the grasses used by local population as fodder [15]. However, no detailed study has carried out to analyze utilization and selection strategies of these plants by shepherds and farmers living in this zone. Extensive areas in the Thal have been overgrazed and they are now strongly threatened by desertification processes [16, 17]. Understanding the relative importance and preference of different species is crucial for a sustainable management of the local forage resources and can help animal husbandry technicians to optimize the selection of useful fodder species and to improve the livestock system efficiency.

Moreover, recording this knowledge would be a much faster and cheaper method for learning about palatability and nutritive value of these plants.

The major aims of this study were:

1. To document traditional knowledge about the use of grasses and sedges as fodder in Thal and to assess similarities and differences with the studies previously conducted in the same [15] and in neighboring areas [11, 12].
2. To evaluate the impact of socioeconomic factors on the local ethnobotanical knowledge
3. To rank, by order of preference, the different species used in the animal diet
4. To quantify the influence of seasonal variation on the availability of these plants as animal feed.

## Materials and Methods

### Area of study

The Thal desert is located between 31° 10’ N and 71° 30’ E in the Punjab province, Pakistan (Fig. 1). It is a subtropical sandy desert lying between the Indus River flood plains in the west and Jhelum and Chenab River flood plains in the east. About 50% of the Thal is under arid to hyper-arid climatic conditions (mean annual rainfall less than 200 mm) and the remaining half is characterized by semiarid climatic conditions (annual mean rainfall between 200 and 500 mm). Most of rainfall occurs between June and August. Average temperatures range between 3-8 °C in winter and 32 – 40 °C in summer. Wind erosion is a serious problem leading to the loss of topsoil and organic matter and damage to crop plants. This region is divided into six districts viz. Bhakkar, Khushab, Mianwali, Jhang, Layyah, and Muzaffargarh.

**Fig. 1:**
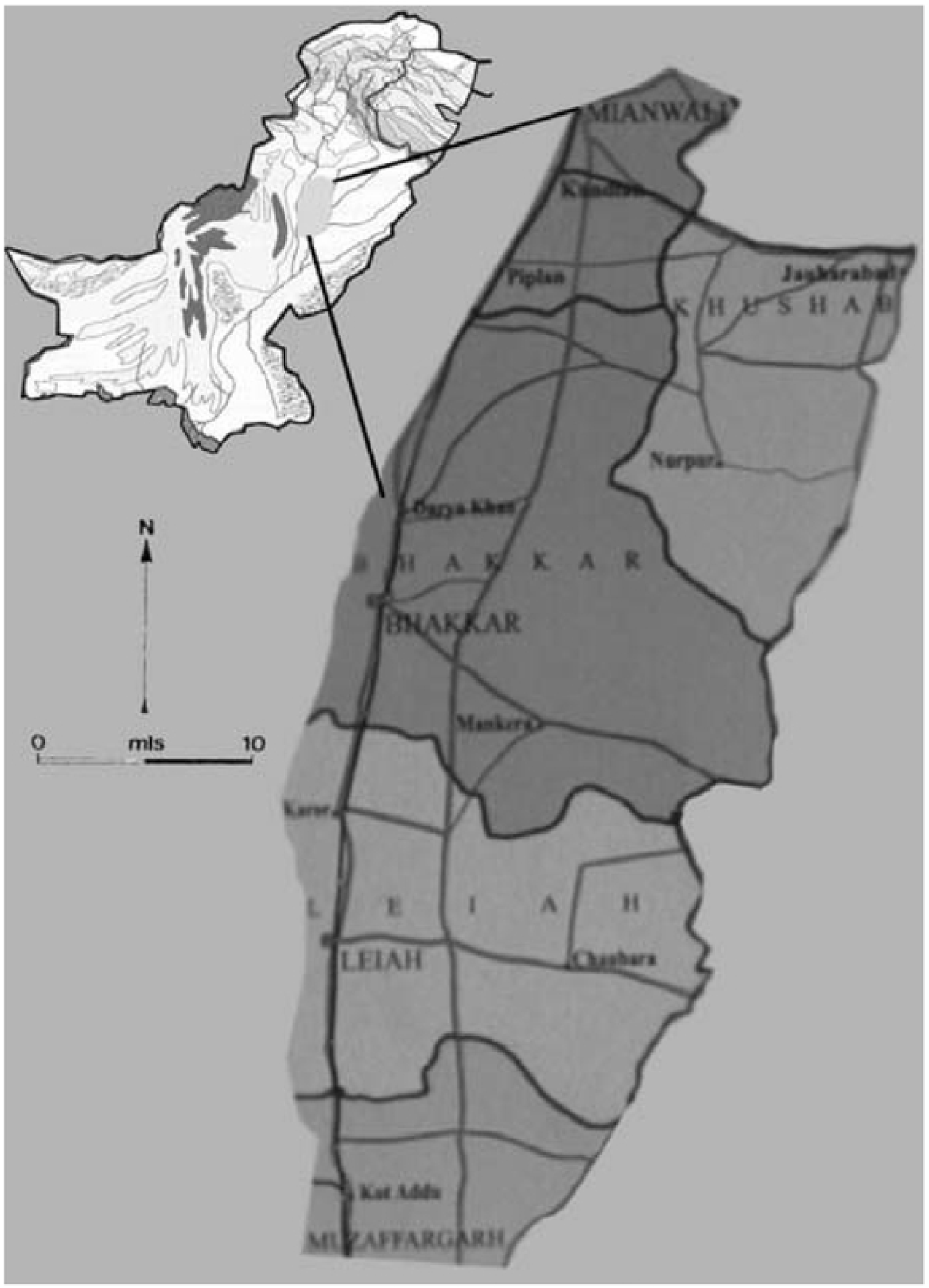
Demography of informants of this study area

In Thal desert livestock is considered as a more secure source of income for small farmers and landless poor people. According to [18] the average herd size is 17 standard animal units. Livestock herds consist of animals of different age and sex; on average each farm has 22.8 goats, 16.7 sheep, 7 cattle, 2.51 buffaloes, 0.88 camels, 0.21 donkeys and 0.05 mules. Detailed information on grazing and stall feeding practiced in the area is given in [19].

### Ethnobotanical survey

Data were collected for two consecutive years (from March 2016 to March 2018) from Thal desert. Thal desert has six districts and we visited each district twice a year for data collection. Informants were selected by snowball-sampling technique [20] among village leaders, shepherds and both farm and domestic livestock caretakers. Formal ethical consent was obtained from all participants before the research started. Information was gathered by using different approaches i.e. group discussions with informants, individual semi-structured questionnaires and participant observation (Fig. 2) [21]. The questionnaires were drafted in the local language (*Seriki* and *Punjabi*) and included the following major questions: (i) Which grasses/sedges are used as fodder? (ii) Which grasses/sedges are the preferred feed of choice for cattle, sheep, camels, buffaloes, and goats? (iii) What is the palatability of the different used plants? (iv) Which plant part do animals consume? (v) What are the feeding habits of different animals? (vi) Which livestock feeding system does local people adopt: free grazing or cut and carry? (vii) Do the listed fodder plants have any ethnoveterinary use? (viii) What are their other indigenous uses?

**Fig. 2:**
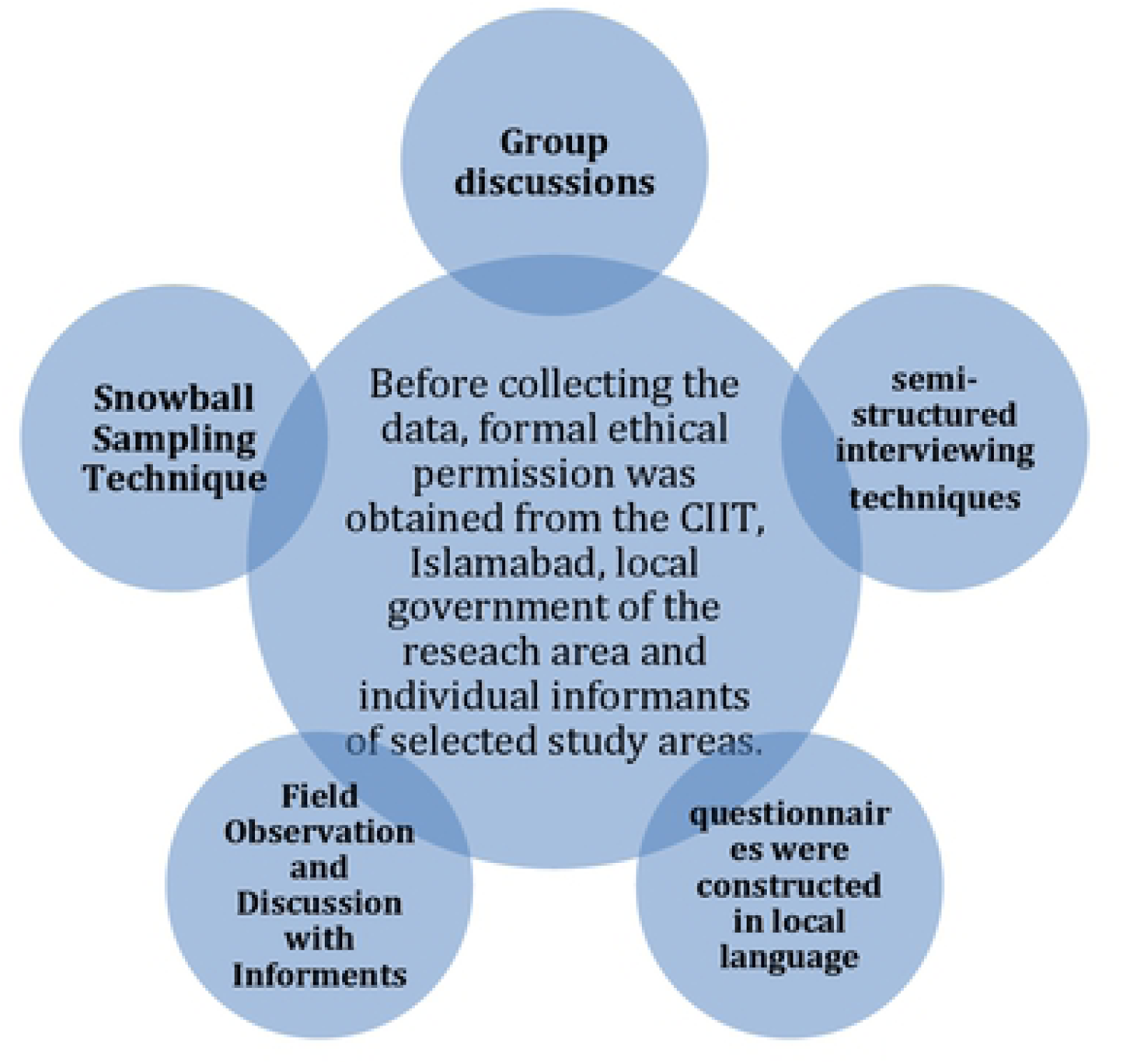
Ethnobotanical survey and data collection

In the second stage of the field research we used direct observation of livestock grazing habits to evaluate the palatability of different plants, animal preferences and the growth stages of plants at the time of grazing.

### Collection and identification of plants

Plant collection was performed with the help of local informers during the field survey. Identification of the gathered species was carried out by the herbarium specialist Dr. Mushtaq Ahmed from Quaid-i-Azam University Islamabad and by the taxonomist Dr. Humaira Shaheen (Fig. 3). Botanical nomenclature of species and families complies with online Flora of Pakistan (http://www.efloras.org/flora_page.aspx?flora_id=5) [22] and the herbarium specimens were kept in to the Botany Department of Pir Mehr Ali Shah University of Arid Agriculture.

**Fig. 3:**
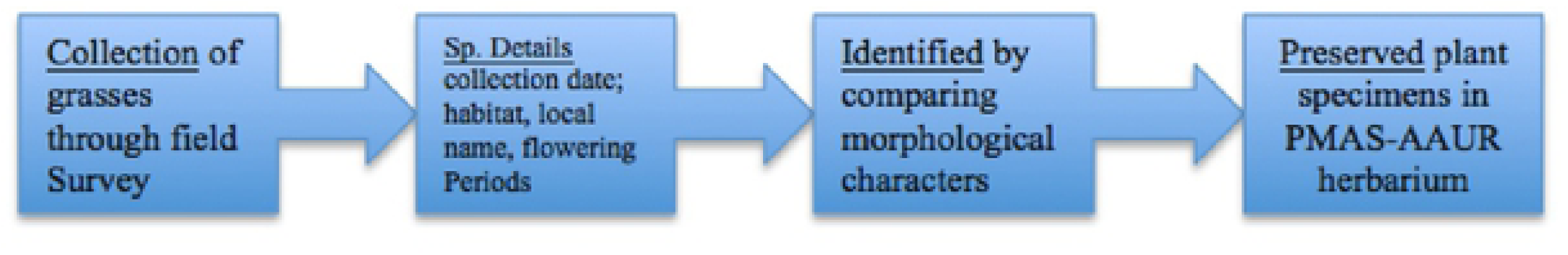
Different steps for collection and identification of Grasses

### Data analysis

The most common method to measure relative abundance was visual assessment and observation of ethnobotanically important grasses in the study area[12]. Total study area was almost 20,000 square kilometers. We randomly divided each district into 45-50 plots and plot size was (10X10m = 100m^2^). Results were constructed by percentage of relative abundance through the following formula;

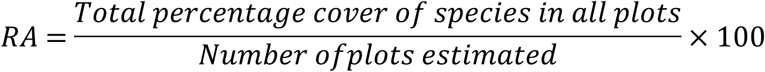

Based on the abundance value, grasses were categorized into the following groups i.e. Abundant, Common, Frequent, Occasional and Rare (Table 1).

**Table 1:** Relative abundance categories and Coverage in the study area

| **Abundance**<br>**scale** | **Abundance**<br>**categories** | **Coverage of**<br>**Grasses** |
| --- | --- | --- |
|  | Rare (R) | <7% |
| 1 | Occasional (O) | 7-10% |
| 2 | Frequent (F) | 10-25% |
| 3 | Common (C) | 25-55% |
| 4 | Abundant (A) | 55-100% |

Relative frequency of citation (RFC) was calculated to sort listed plants by priority order, using the following formula [23–25].

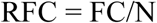

Where FC is the number of informants that mentioned the fodder use of the species and N is the total number of informants included in the study.

Pairwise comparison (PWC) was also used to determine the priority order of the listed species [12]. Ten informants (5 key informants and 5 randomly selected) were chosen for the PWC. The participants were asked, one at a time, to select their preferred fodder plants from all possible pairs of species. Each species got a score of 1 if the participants selected it. The final score was obtained by adding the scores and ranking them.

Smith’s salience index and Composite Salience [26] were used to judge species saliency by weighing the average of the inverse rank of a species across multiple free-lists where each list was weighed by the number of species in the list. ANTHROPAC [27] was used to generate Smith’s salience indexes.

Pairwise ranking or comparison was used to evaluate the degree of preference or levels of importance. The values for use reports across the selected species were summed up and ranked. Ten informants (six key and four randomly taken informants) in the study area ranked grasses according to their use e.g. 1st, 2nd, 3rd, 4th and 5^th^ respectively. Ranking can be used for evaluating the degree of preference or level of importance of selected plants [26, 28].

### Socioeconomic factors

In total, 232 local informants were interviewed (Table 2); 141 were men and 91 were women. A smaller number of female informants was expected and can be partially explained with the local cultural restrictions preventing women from working outside their homes or farms. Informants were grouped into three major age categories: 20–35 (48 informants), 36–50 (116 informants) and 51–67 years (68 informants). With regard to the profession, 34% (36 females and 44 males) were shepherds, 26% (27 females and 33 males) were farmed livestock caretakers and 40% (28 females and 64 males) domestic livestock caretakers. Thirty-six (16%) of the interviewed people were illiterate, 24 (10%) never completed their primary education, 120 (52%) completed 5 years of primary school and 52 (22%) informants had middle education level (Fig. 4) [22].

**Fig. 4:**
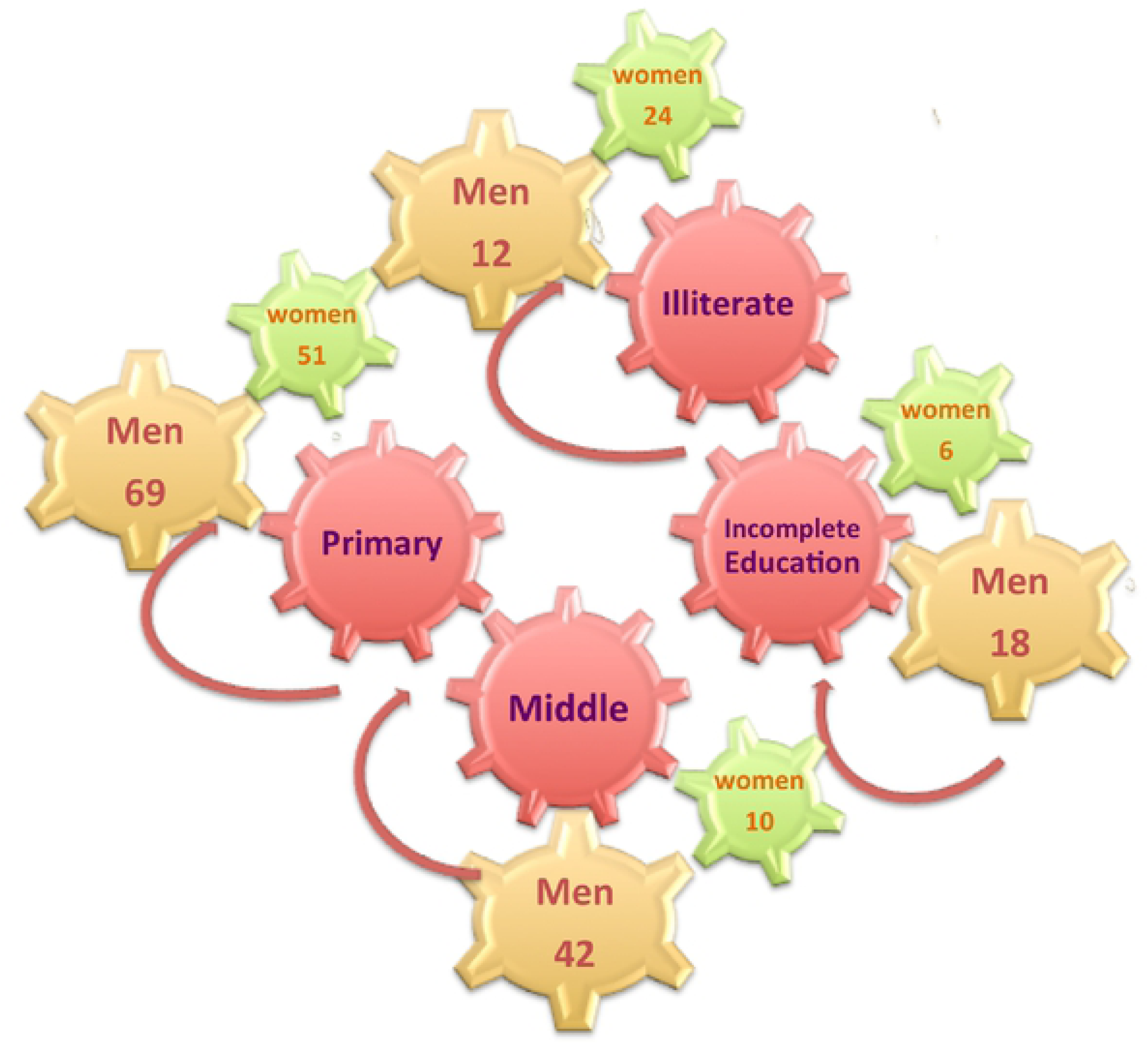
Summary of education levels of informants

**Table 2:** Demography of informants of the study area

| Type of Informants | Young<br>aged<br>20-35 | Middle<br>aged<br>36-50 | Seniors<br>aged<br>51-67 | Total |
| --- | --- | --- | --- | --- |
| Local Shepherds (F) | 8 | 19 | 9 | 36 |
| Local Shepherds (M) | 11 | 20 | 13 | 44 |
| Farmed Ruminant care takers (F) | 5 | 17 | 5 | 27 |
| Farmed Ruminant care takers (M) | 11 | 16 | 6 | 33 |
| Domestic Ruminant care takers (F) | 7 | 12 | 9 | 28 |
| Domestic Ruminant care takers (M) | 6 | 32 | 26 | 64 |
| Total informants | 48 | 116 | 68 | 232 |

**Key:** Local Shepherds (who take care cattle in the field for free grazing), Farmed Ruminant caretakers (who take care cattle in the livestock forms), Domestic Ruminant caretakers (who take care cattle in their home).

## RESULTS AND DISCUSSION

### Use of fodder species

The informants reported the use of 61 plant species that were distributed into 40 genera and 3 botanical families. The most represented genus was *Cyperus* with 5 species, followed by *Cenchrus* and *Eragrostis* with 4 species each. Most species belonged to Poaceae family (51 species; 84% of the reported plants) while 8 species (13%) were categorized into Cyperaceae family. Typhaceae were represented by only one species: *Typha elephantina*. Fifty-five species (92% of the reported species) were classified as native and 5 (8%) as exotic. The following exotic species were reported by informants: *Chloris gayana*, *Imperata cylindrica*, *Paspalum dilatatum*, *Sorghum bicolor* and *Vetiveria zizanioides.* These results seem to reflect composition and distribution patterns of the local flora. In a floristic checklist of Thal desert, Shaheen et al. [22] observed that Poaceae was the leading family with 52 species. Of the 52 Poaceae naturally occurring in the area, 48 (94%) were reported to be used as fodder in our study; 5 were not cited by informants and 4 (*Brachiaria reptans*, *Eragrostis atrovirens*, *E. cilianensis*, *Themeda anathera*) were reported in our study but not in the floristic inventory. All the eight Cyperaceae cited were included in the study conducted by Shaheen et al. [14].

Our comparative analysis revealed 15 species that are used as fodder in all the considered studies. We found a mean similarity (Jaccard’s index) rather high (36.4 ± 6.9) with values ranging from 30.8 (this study vs [11]) to 50.0 ([12] vs [11]). These studies are all from zones lying in the proximity of the study area that share not only similar ecological factors but also the same socioeconomic and cultural history. Nevertheless, our study listed 20 grasses not previously reported for this area in the fodder category. These results provide an important contribution of novelty to the knowledge on wild fodder plants in Pakistan. At the same time, they also show the importance of collecting new ethnobotanical information even in areas already studied.

### Socioeconomic factors

Informants mentioned 8.27 ± 4.49 taxa (range 1 – 18). Gender (H = 0.373; P > 0.05) and education (H = 5.29; P> 0.05) had no influence on the knowledge of fodder plants. Gender influence on traditional knowledge is controversial [29] and many studies have showed that the statistical strength of this relation depends on the local cultural context and on the categories of use that the researchers focus on. A lack of differentiation between men and women, as observed in this study, could mean that there is not a clear division of labor in the area. A similar finding was observed by Aumeeruddy et al. [30] in Northern Pakistan, where women have a detailed knowledge on characteristics and properties of the different fodder species, suggesting that they fully share with men the responsibility of livestock rearing and forage collection. Khan and Khan [31] observed that most of the women of Cholistan desert have an important role in managing livestock, spending almost 8 to 13 hours a day in this activity. Differently Nunes et al. [7] and Bruschi et al. [6] showed that men prevail in the knowledge about fodder plants. The greater male knowledge found in these two studies may be explained by different gender-based experiences and skills: men spend much of their time moving with their herds while women are more frequently involved in managing food and family care. The age of informants resulted to be statistically significant (H = 9.97; P < 0.05). As also shown in many other ethnobotanical studies [32]; [33]; [34], elderly people seem to retain more traditional knowledge on the use of plants. For young people (25 – 35 years old), the average number of known fodder plants was 6.65 ± 4.12 while for middle-aged (36 – 50) and elderly informants (> 50) there was an average number of 8.25 ± 4.13 and 9.42 ± 4.74, respectively. Occupation also strongly affected the number of fodder species reported by informants (H = 14.58; P < 0.01). Domestic livestock caretakers mentioned a higher number of plants (9.50 ± 4.43) followed by farmed livestock caretakers (7.98 ± 4.02) and shepherds (7.10 ± 4.60). Domestic livestock caretakers spend much time with cattle; therefore have a better knowledge about the animals’ favorite foods.

### Pairwise ranking of wild palatable plants

*Cymbopogon jwarancusa* subsp. *jwarancusa* with 1^st^ rank was the most preferred species among all selected grass species, followed by *Cynodon dactylon, Cenchrus ciliaris, Typha elephantina* and *Cyperus alopecuroides* which had 2^nd^ 3^rd^, 4^th^ and 5^th^ rank respectively. *Pycreus flavidus* received the lowest score, therefore resulting as the less preferred species (Table 3). The most highly ranked species (*Cymbopogon jwarancusa* subsp. *Jwarancusa, Cynodon dactylon, Cenchrus ciliaris, Typha elephantina* and *Cyperus alopecuroides*) are also the most dominant in the area (Shaheen, unpublished data). This finding seems to support the “appearance hypothesis” under which the most abundant species are better known and mostly used Lucena et al. [35]. Plants commonly growing in given area would allow local people to have more experience of their properties and consequently would have a greater probability of being introduced into the local culture.

**Table 3:** Pair wise ranking of wild palatable plants from all districts of Thal

| S. No. | Botanical name | R1 | R2 | R3 | R4 | R5 | R6 | R7 | R8 | R9 | R10 | T | R |
| --- | --- | --- | --- | --- | --- | --- | --- | --- | --- | --- | --- | --- | --- |
| 1 | *Cymbopogon jwarancusa* subsp. *jwarancusa* (Jones) Schult. | 5 | 5 | 5 | 5 | 5 | 5 | 5 | 5 | 5 | 3 | 48 | 1<sup>ST</sup> |
| 2 | *Cynodon dactylon* (L.) Pers. | 5 | 4 | 4 | 5 | 4 | 4 | 4 | 5 | 4 | 4 | 43 | 2<sup>ND</sup> |
| 3 | *Cenchrus ciliaris* L. | 4 | 3 | 4 | 4 | 4 | 5 | 3 | 4 | 4 | 4 | 39 | 3<sup>RD</sup> |
| 4 | *Typha elephantina* Roxb. | 5 | 4 | 5 | 3 | 3 | 5 | 4 | 5 | 3 | 1 | 38 | 4<sup>TH</sup> |
| 5 | *Cyperus alopecuroides* Rottb. | 4 | 2 | 3 | 3 | 4 | 3 | 5 | 4 | 2 | 3 | 33 | 5<sup>TH</sup> |
| 6 | *Eragrostis minor* Host | 2 | 2 | 3 | 4 | 4 | 5 | 2 | 2 | 3 | 5 | 32 | 6<sup>TH</sup> |
| 7 | *Sporobolus arabicus* Boiss. | 2 | 3 | 4 | 4 | 3 | 2 | 3 | 2 | 3 | 5 | 31 | 7<sup>TH</sup> |
| 8 | *Brachiaria reptans* (L.) C. A. Gardner & C.E. | 1 | 5 | 4 | 2 | 3 | 1 | 0 | 4 | 5 | 5 | 30 | 8<sup>TH</sup> |
| 9 | *Tragus roxburghii* Panigrahi | 1 | 5 | 4 | 2 | 3 | 1 | 0 | 4 | 5 | 5 | 30 | 9<sup>TH</sup> |
| 10 | *Lasiurus indicus* Henr. | 4 | 2 | 2 | 4 | 5 | 3 | 2 | 2 | 4 | 1 | 29 | 10<sup>TH</sup> |
| 11 | *Aristida funiculata* Trin. & Pupr. | 5 | 4 | 2 | 3 | 1 | 0 | 4 | 5 | 3 | 2 | 29 | 10<sup>TH</sup> |
| 12 | *Cenchrus pennisetiformis* Hochst. & Steud. | 1 | 5 | 4 | 2 | 3 | 1 | 0 | 4 | 5 | 4 | 29 | 10<sup>TH</sup> |
| 13 | *Saccharum spontaneum* L. | 2 | 2 | 3 | 4 | 4 | 5 | 2 | 2 | 3 | 2 | 29 | 10<sup>TH</sup> |
| 14 | *Themeda triandra* Forsk. | 5 | 4 | 2 | 3 | 1 | 0 | 4 | 5 | 3 | 2 | 29 | 10<sup>TH</sup> |
| 15 | *Pycreus flavidus* (Retz.) T. Koyama | 2 | 3 | 3 | 2 | 4 | 1 | 3 | 2 | 3 | 5 | 28 | 11<sup>TH</sup> |

### Co-relation used for pairwise comparison

On the basis of RFC value, pairwise comparison was used to correlate fodder grasses and the knowledge of the respondent. Ten out of 232 respondents were chosen on the basis of their profession (ethnoveterinary practitioner) but were potential respondents due to enough indigenous knowledge. Based on RFC values knowledge of respondent R1 showed a strong correlation with R4, as R2 (0.56; p<0.001) showed with R1 with R7 (0.55;p<0.001), R2 have strong correlation with R3 and R8 (0.48, 0.58; p<0.001) but R2 had the strongest correlation with R9 (0.71; p<0.001). All correlation and their distribution of RFC values are shown in Fig. 5. The positive correlation between respondents suggests that respondents report similar information about the plant, for example, R2 and R9 both were an ethnoveterinary practitioner with age more than 50 so they have similar knowledge.

**Fig. 5:**
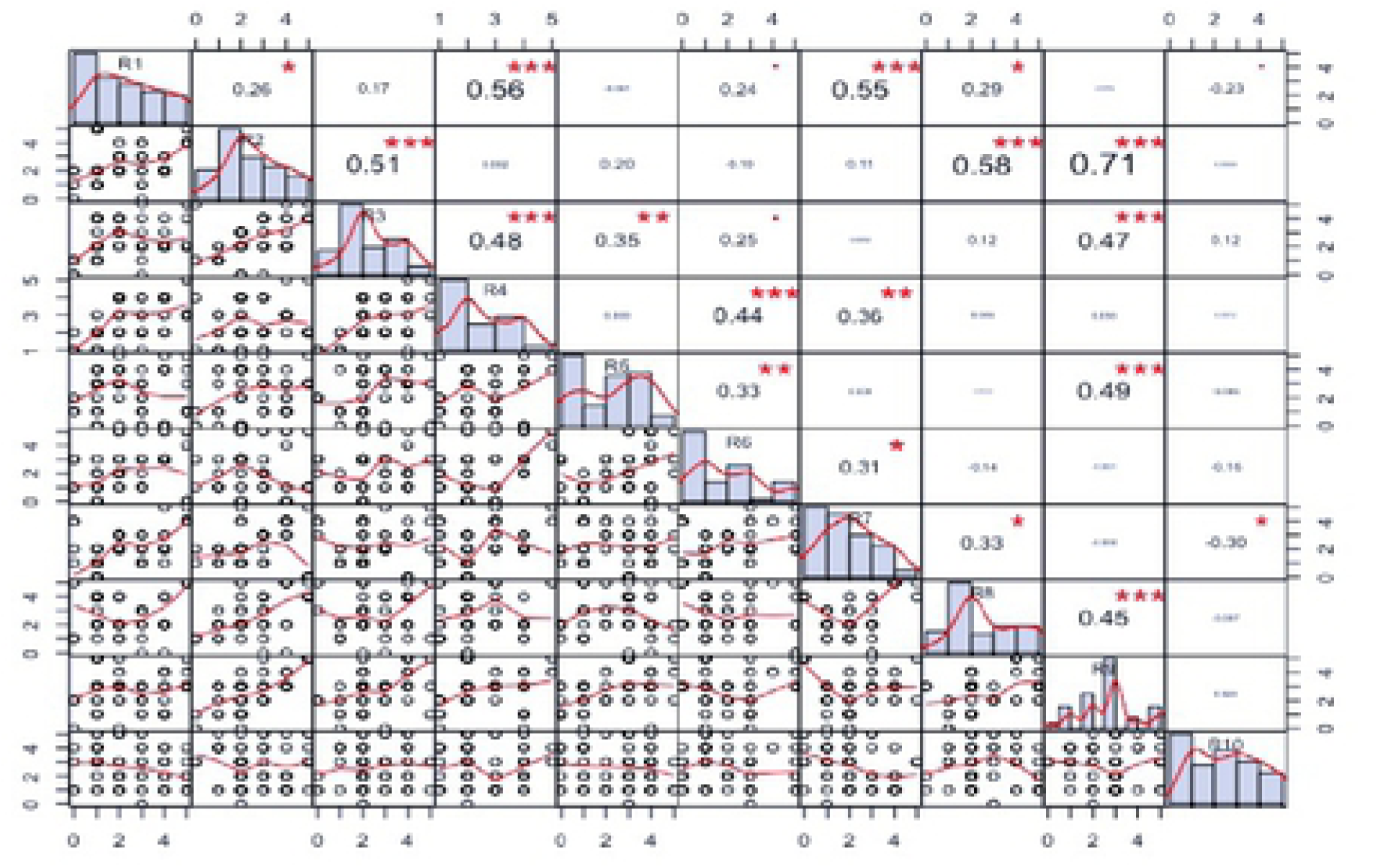
Co-relation used for pairwise comparison for different grasses

### Availability and Prioritizing fodder grasses on the basis of RFC and PWC

RCF values ranged from 1 to 0.51 with a mean value of 0.71. Twenty-five species had RFC values higher than average value while the remaining 35 species had RFC value lower than the average value (Fig. 6, Table 4). *Cymbopogon jwarancusa* and *Cynodon dactylon* showed the highest value (1.00) while *Imperata cylindrical* (0.52) and *Vetiveria zizanioides* (0.51) had the lowest values. Based on these RFC values fodder species were included into three categories of priority: species with higher priority (group A), species with medium priority (group B) and species with low priority (group C). Twenty-eight (45.9%) species were highly preferred by the informants followed by twenty-three (37.7%) species that had medium priority while ten (16.3%) grass species were the least preferred (Fig 7). Values ranged between 1-0.69 for group A, between 0.69-0.54 for group B and between 0.54-0.51 for group C. Similar results were shown by Harun et al. [12] in their study. These results were confirmed by cluster analysis based on RFC in which the reported species were classified into three major groups compliant with the results of priority ranking analysis. Similar results were found when we performed cluster analysis using PWC data. *Cymbopogon jwarancusa* was the preferred species in both approaches (Table 5).

**Fig. 6.**
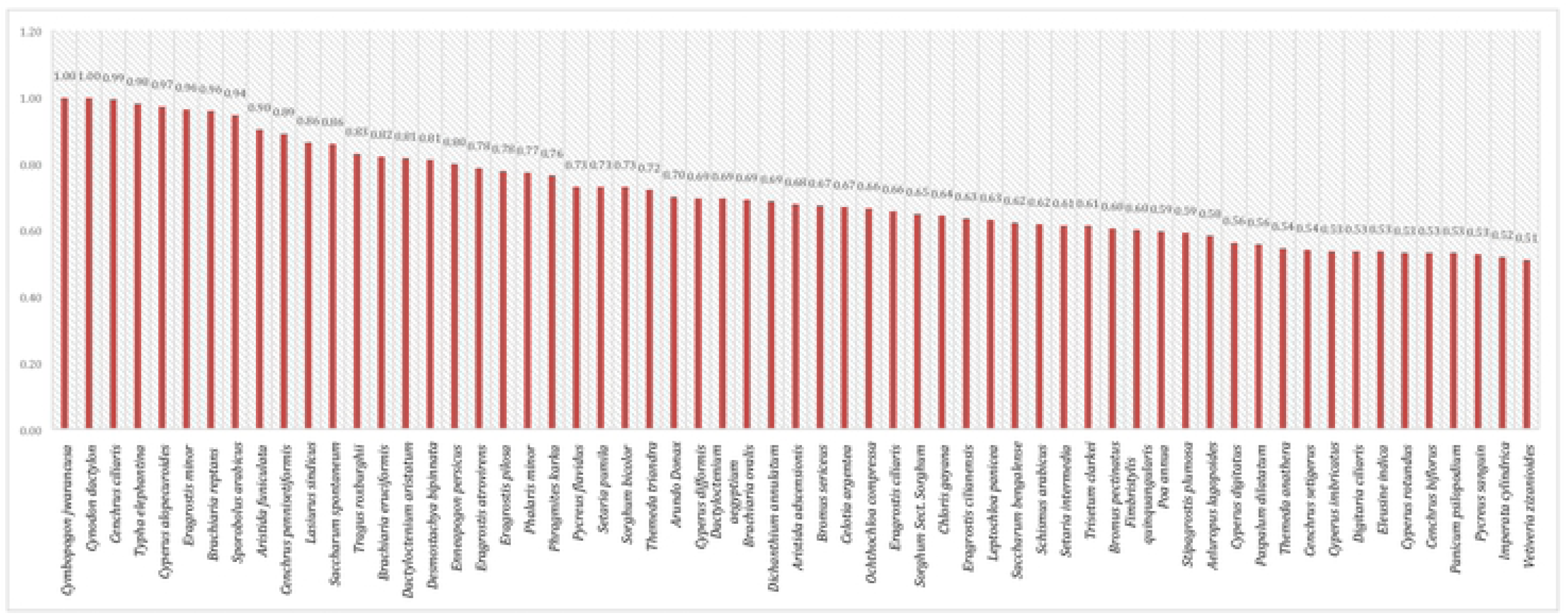
Prioritizing of fodder grasses on the bases of RFC

**Fig. 7:**
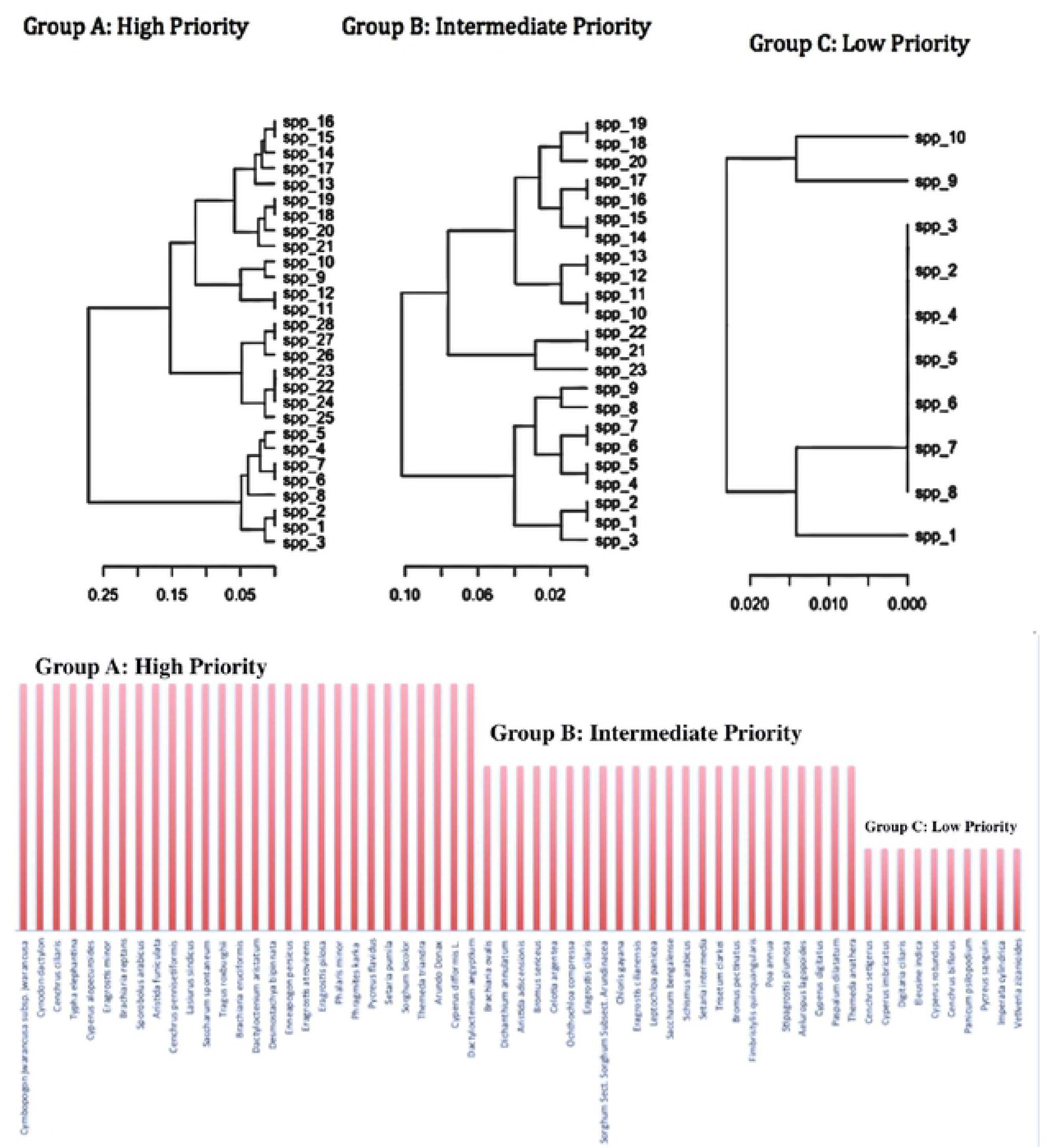
Cluster analysis for grouping of ethno botanically used fodder grasses

**Table 4:**
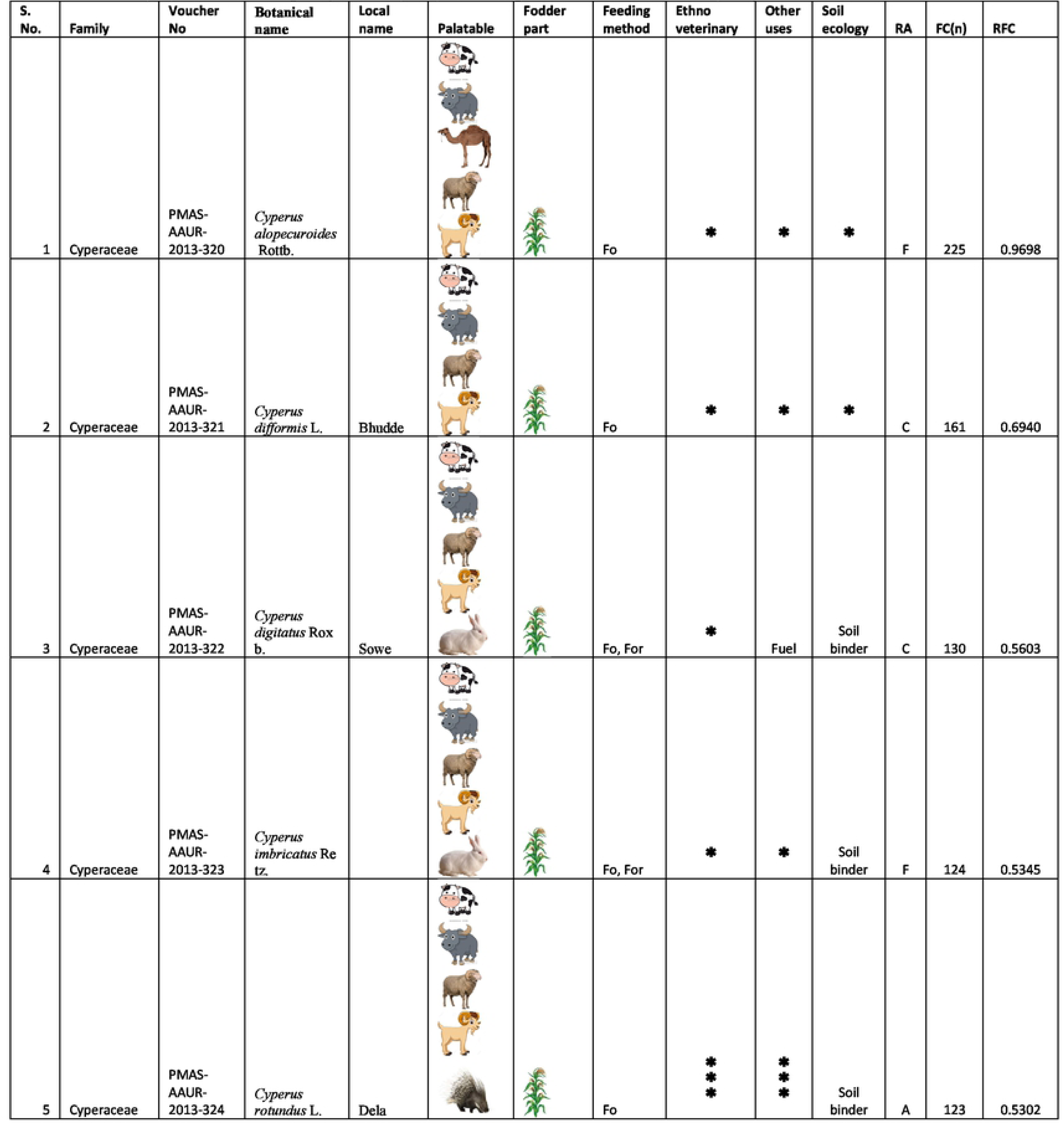
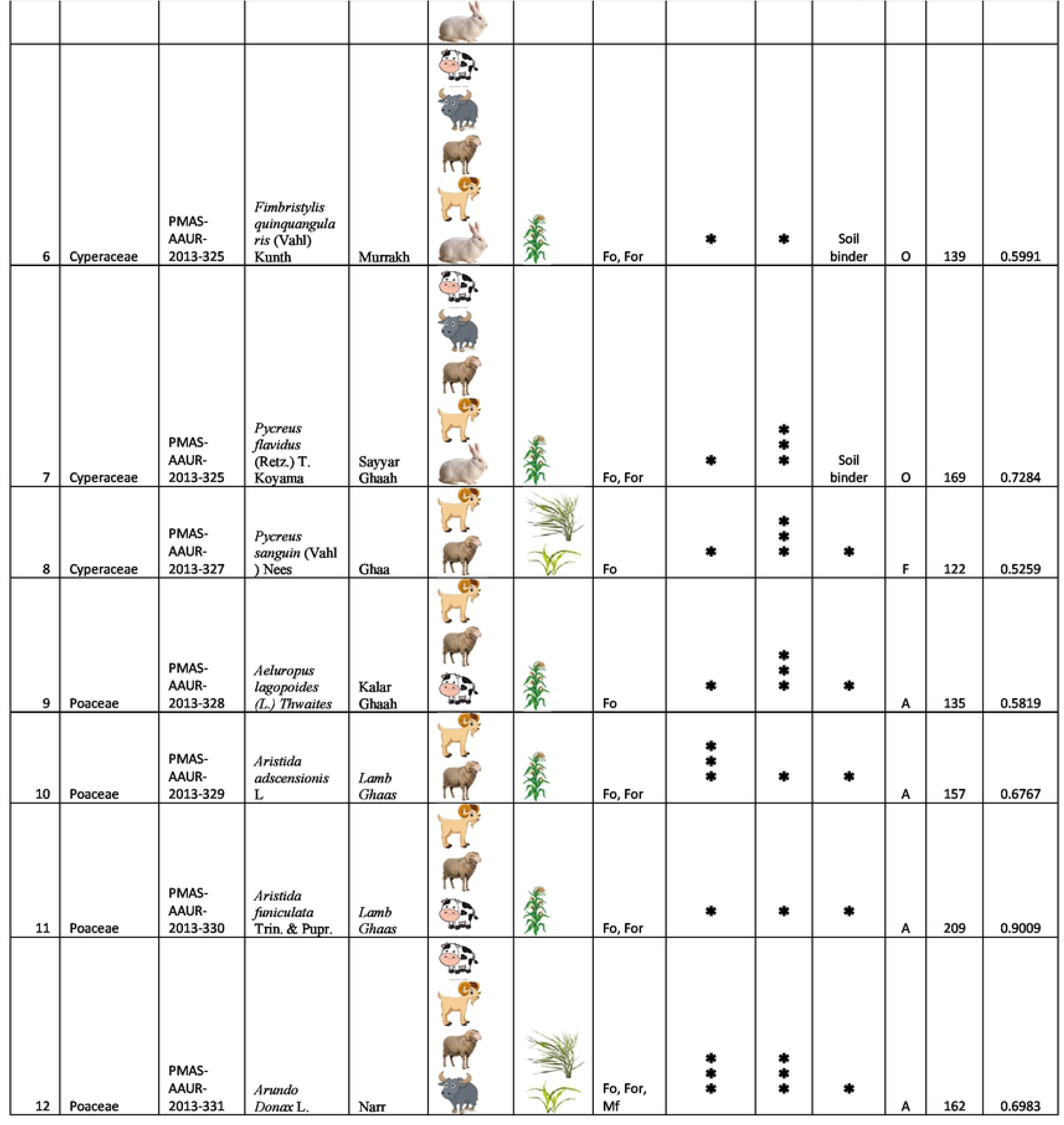
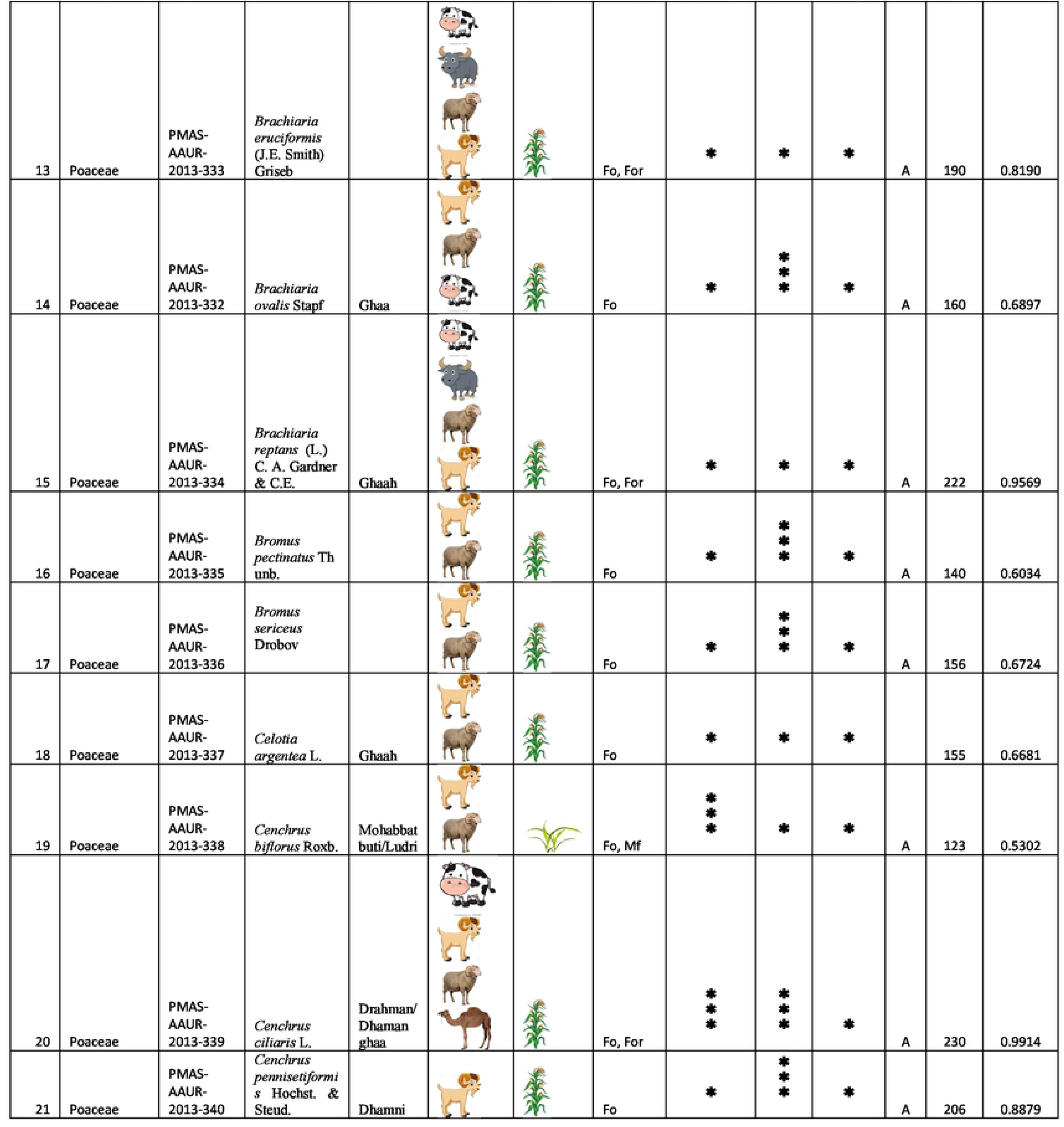
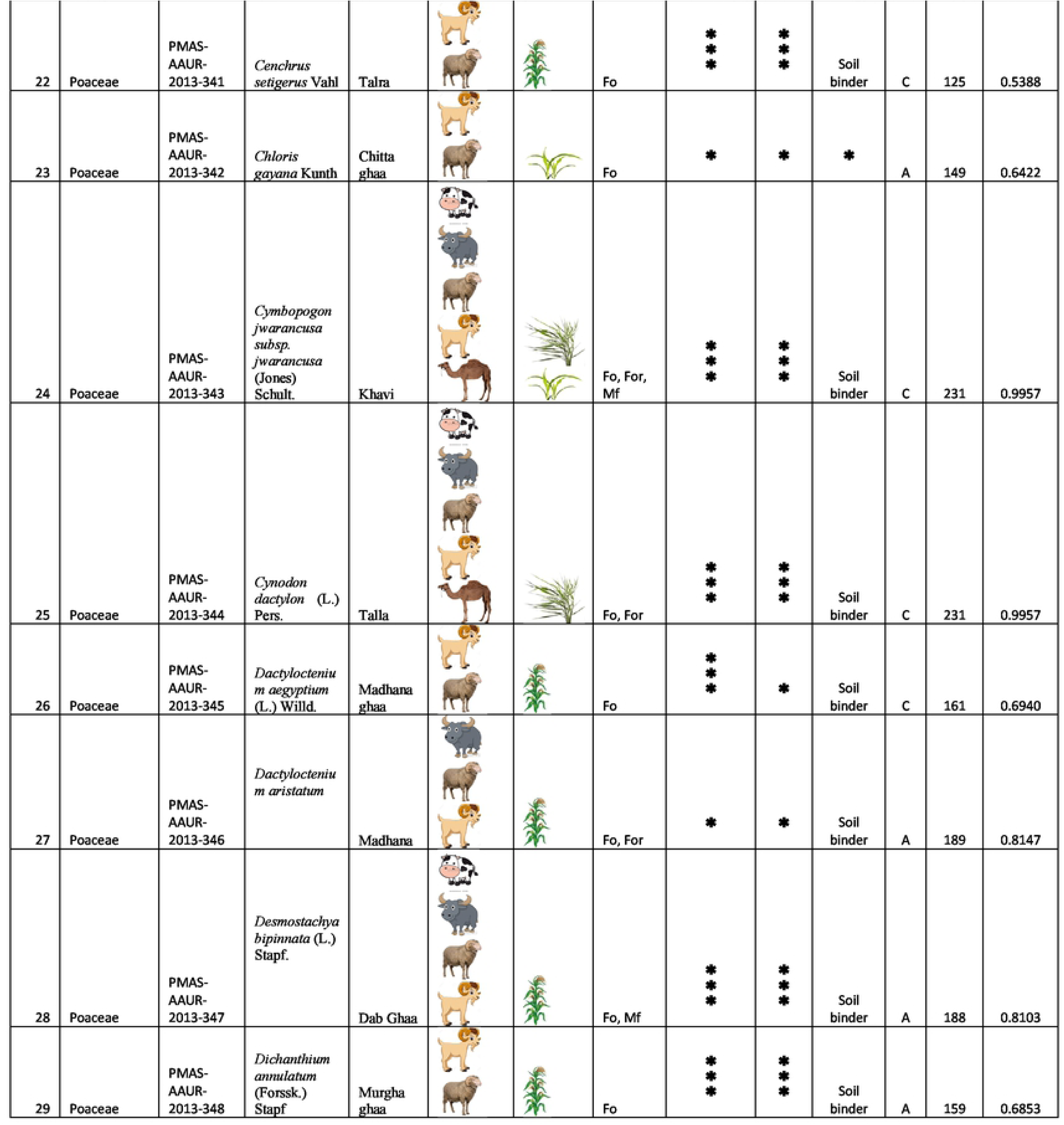
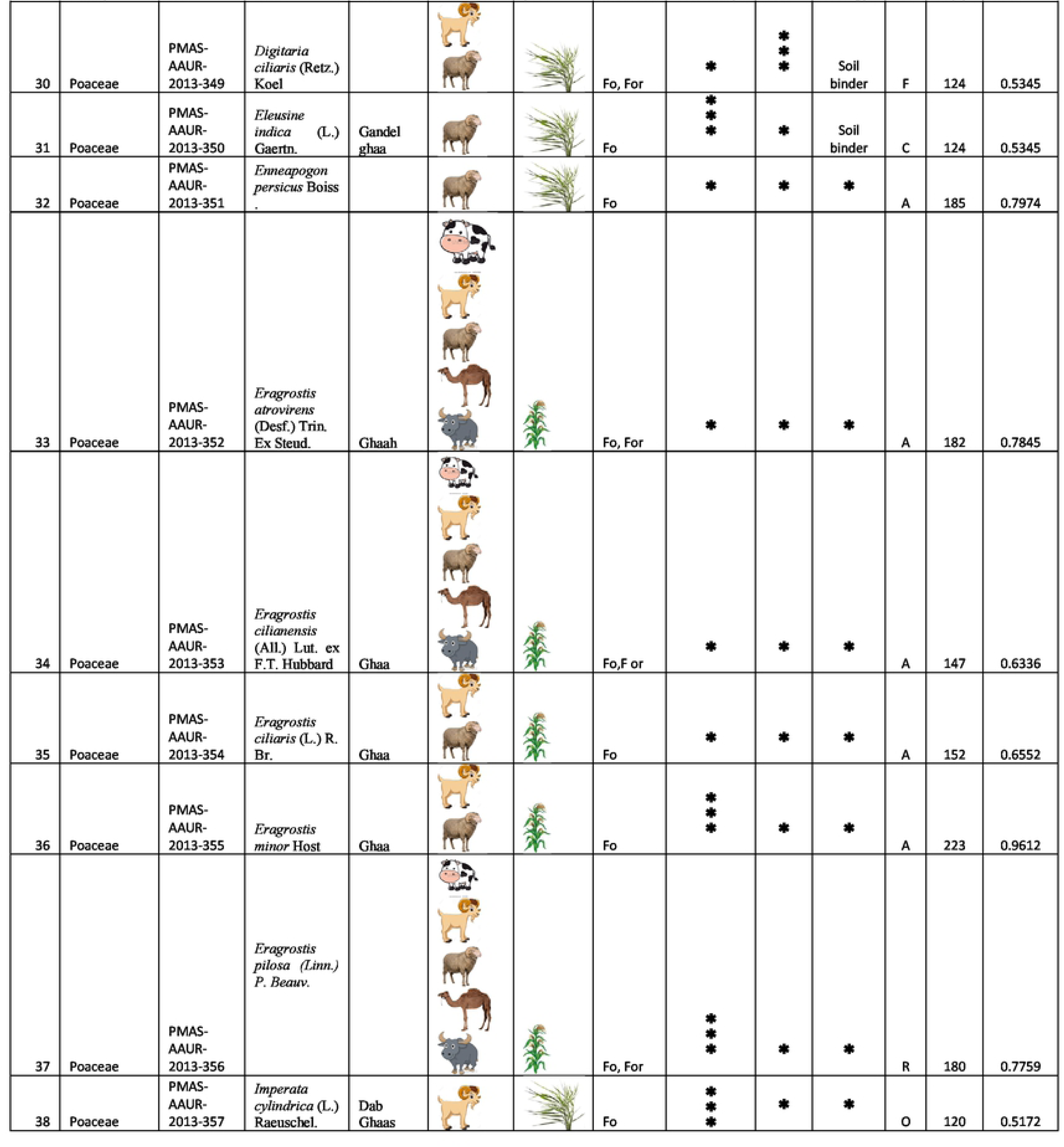
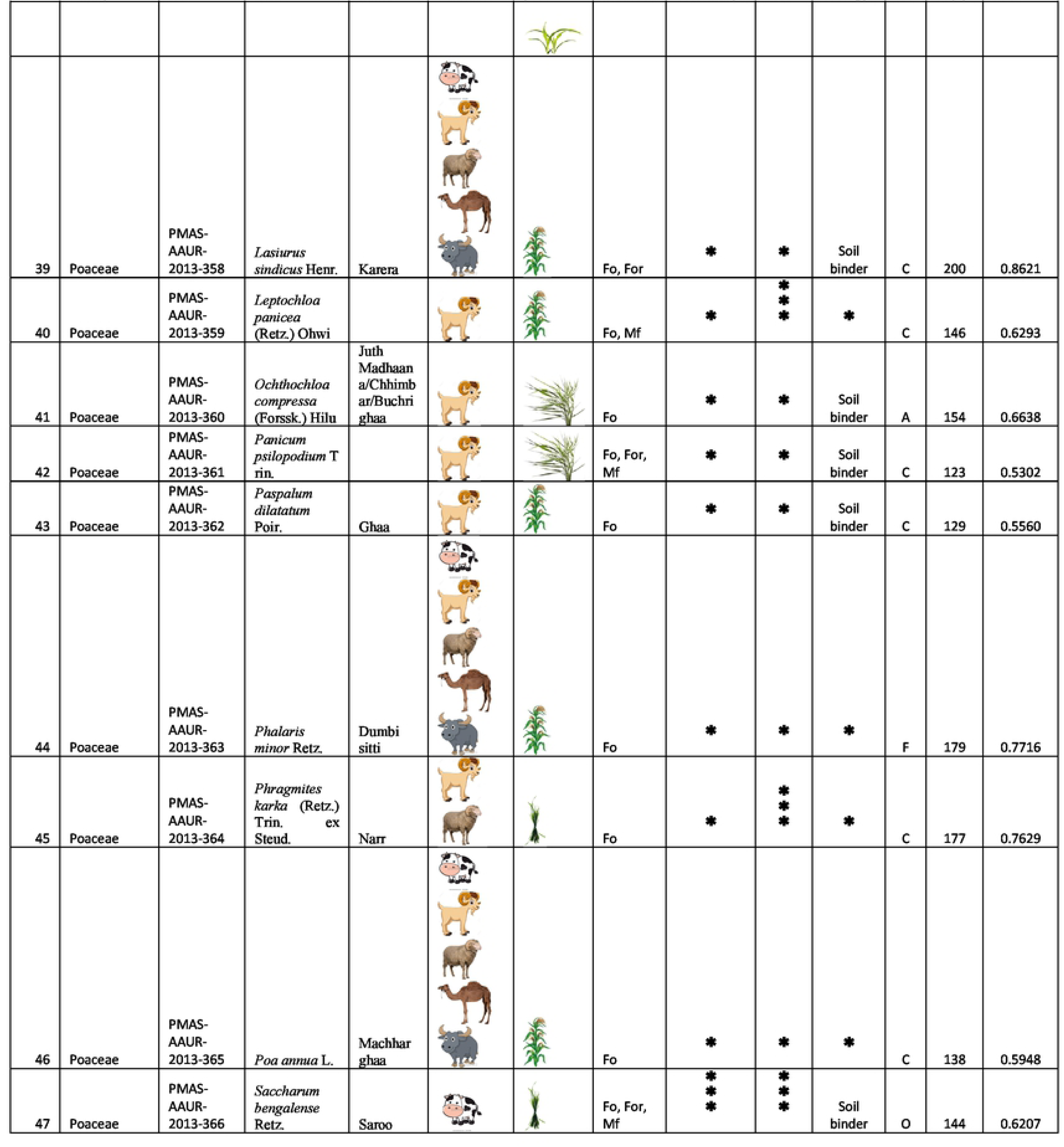
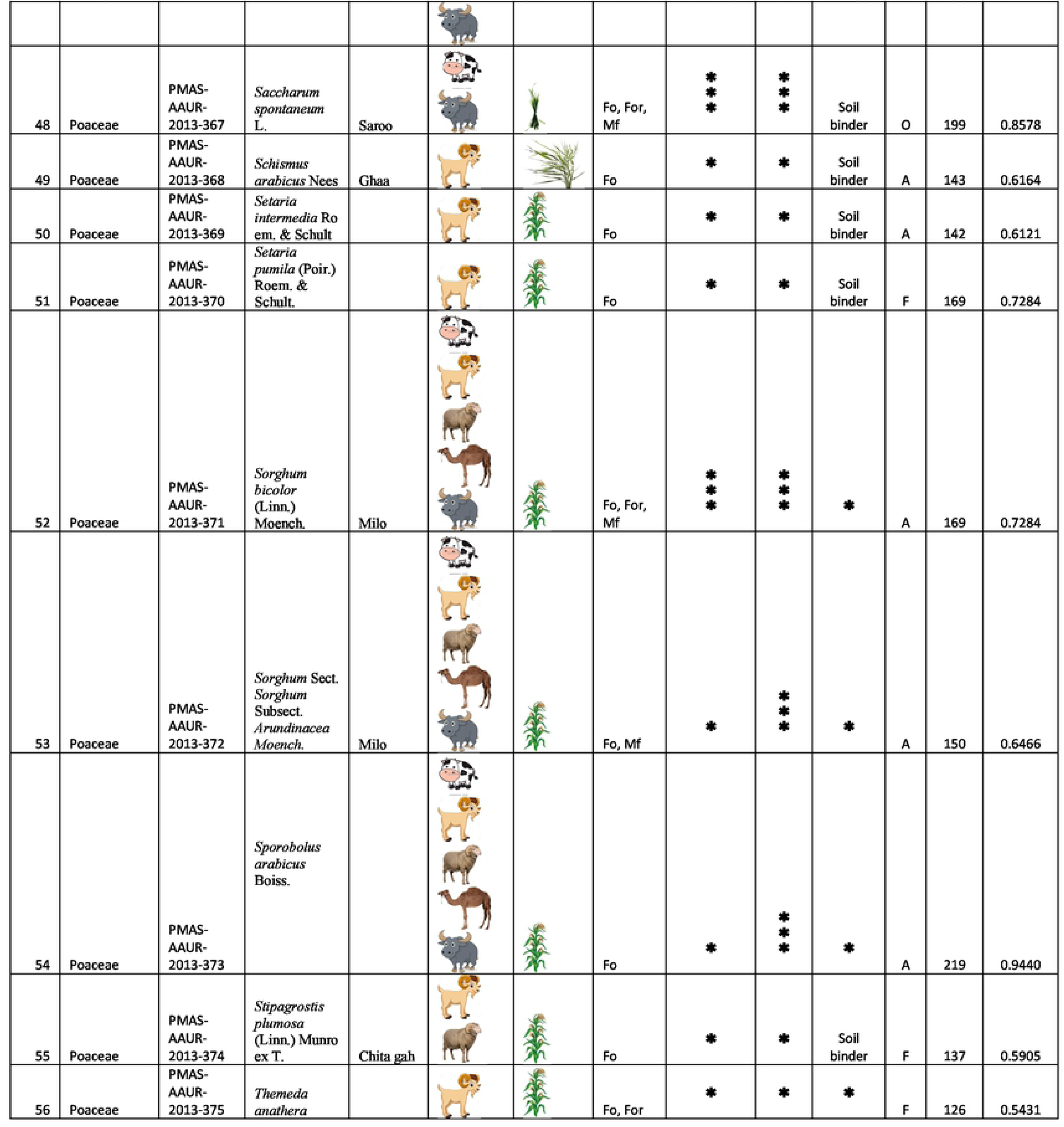
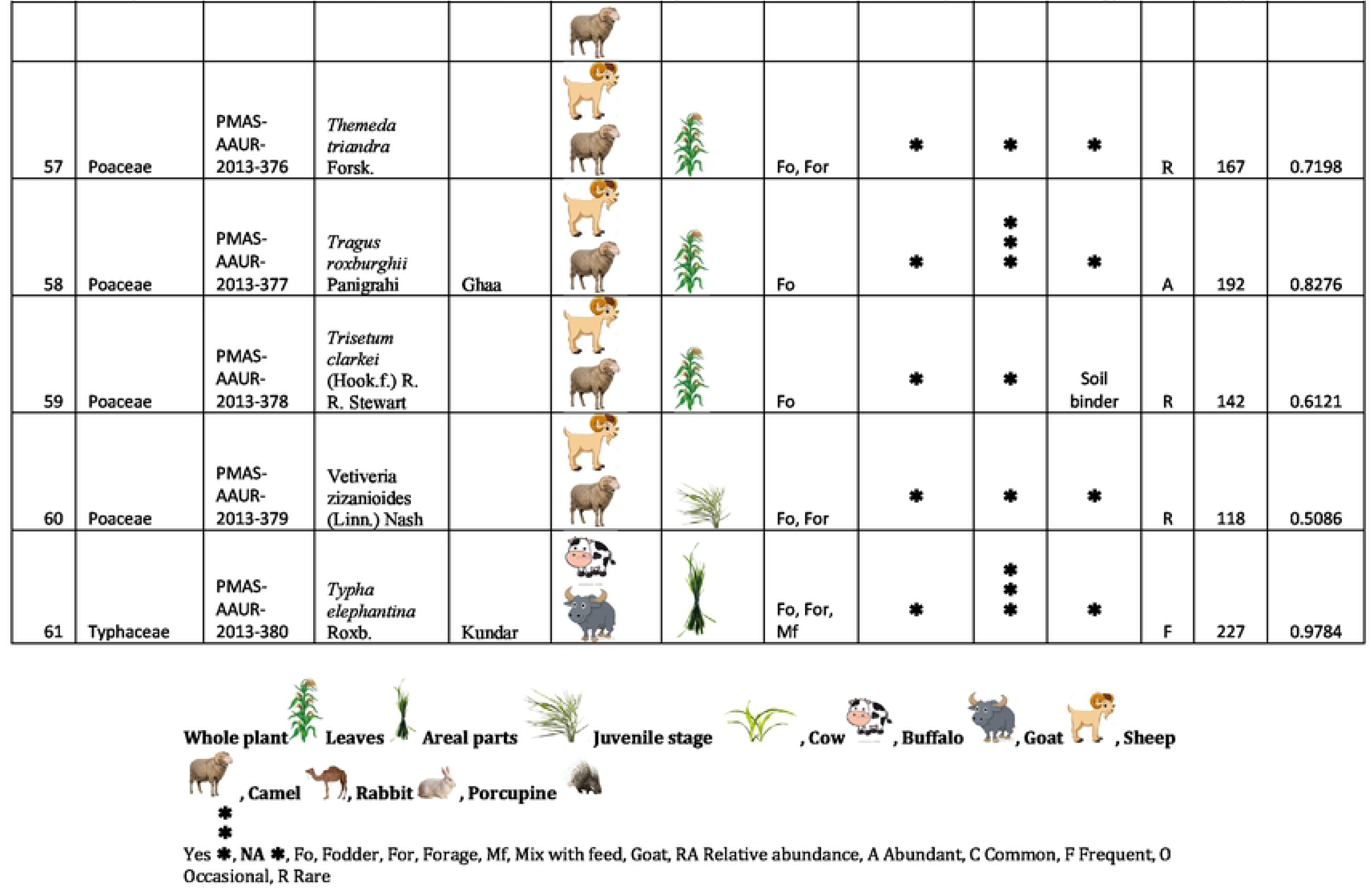
List of the collected grasses, Ethnobotanical, Ethno veterinary data, abundance; focal persons count (FC) and relative frequency citation (RFC) of fodder grasses of area of Thal desert, Punjab Pakistan

**Table 5:** Pairwise comparison (PWC) base on similar RFC vales of fodder grasses

| Fodder grasses | Total gained % points | Rank |
| --- | --- | --- |
| **GROUP A (RFC =0.9957-0.9009)** |  |  |
| *Cymbopogon jwarancusa subsp. jwarancusa* | 88.2 | 1st |
| *Typha elephantina* | 87.3 | 2nd |
| *Cynodon dactylon* | 87.1 | 3rd |
| *Cenchrus ciliaris* | 85.1 | 4th |
| *Cyperus alopecuroides* | 84 | 5th |
| **GROUP B (RFC =0.8879-0.8103)** |  |  |
| *Cenchrus pennisetiformis* | 72.5 | 1st |
| *Lasiurus indicus* | 63.5 | 2nd |
| *Saccharum spontaneum* | 62.4 | 3rd |
| *Tragus roxburghii* | 60.9 | 4th |
| **GROUP C (RFC =0.7974-0.6940)** |  |  |
| *Enneapogon persicus* | 77.9 | 1st |
| *Eragrostis atrovirens* | 76.8 | 2nd |
| *Eragrostis pilosa* | 72.1 | 3rd |
| *Phalaris minor* | 70.1 | 4th |
| *Phragmites karka* | 61.1 | 5th |
| **GROUP D (RFC =0.6897-0.6121)** |  |  |
| *Brachiaria ovalis* | 72.1 | 1st |
| *Dichanthium annulatum* | 60.3 | 2nd |
| *Aristida adscensionis* | 59.9 | 3rd |
| *Bromus sericeus* | 58.7 | 4th |
| *Celotia argentea* | 55.9 | 5th |
| **GROUP E (RFC =0.6034-0.6)** |  |  |
| *Bromus pectinatus* | 92.8 | 1st |
| *Fimbristylis quinquangularis* | 90.5 | 2nd |
| *Poa annua* | 85.2 | 3rd |
| *Stipagrostis plumosa* | 76.9 | 4th |
| **GROUP F (RFC =0.5431-0.5086)** |  |  |
| *Themeda anathera* | 59.1 | 1st |
| *Cenchrus setigerus* | 55.6 | 2nd |
| *Cyperus imbricatus* | 54.9 | 3rd |
| *Digitaria ciliaris* | 52.3 | 4th |

The species included in the Group A (high priority) is ecologically dominant and largely available in the area. Moreover, taxa included in this group have a good palatability and are also available during the dry season when other grazing resources are exhausted.

### Palatability of grasses and the method of feeding

Preferred palatability species are often leafy and without lots of stem, with a high leaf table and leaves of low tensile strength [36, 37]. Palatability analysis showed that 77% of the reported species are grazed in the study area (Table 6). In particular; grasses included in group A of the priority ranking were consumed by all ruminants locally raised. Goats are the only animals to feed on every type of grass growing in Thal desert although palatability results show a preference for 58% of the reported species. 40% of the species represented the favorite fodder for sheep and 26% the favorite fodder for buffaloes Camels are very selective animals and use only few specific grasses as fodder (Fig. 8). Different parts showed to have different edibility: for example 42% of grass species were consumed as whole plant (e.g. *Cynodon dactylon*, *Eragrostis minor*, *Cenchrus ciliaris*, *Cenchrus pennisetiformis* etc) while 38% and 19% of them were consumed as aerial parts and as leaves, respectively. The reason why so many grasses are grazed as a whole is probably related to their small size and tender herbaceous texture (e.g. *Cynodon dactylon, Lasiurus sindicus, Phalaris minor, Cyperus rotundus, Eragrostis minor* etc similer results shown in other literature [12] [13]. Due to the sandy nature of soils occurring in the study area these plants have shallow root systems and can easily be pulled out from the soil. Species growing in the form of dense patch are hard to be consumed as a whole plant and animals can enjoy feeding only with the aerial parts of this grass. Beliefs on the feeding habit of the livestock are common in the area: for example, some local shepherds reported that putting the herd out to pasture in open field improves their health and milk production. According to them animals freely grazing are able to select the better grasses avoiding the toxic or less nutritious ones. They justify their belief by comparing milk production of freely grazing animals with cattle fed with forage and also by saying that during dry season, when free grazing is not possible, there is a considerable reduction in animal health and milk production. As Provenza et al describe in their study [38].

**Fig. 8:**
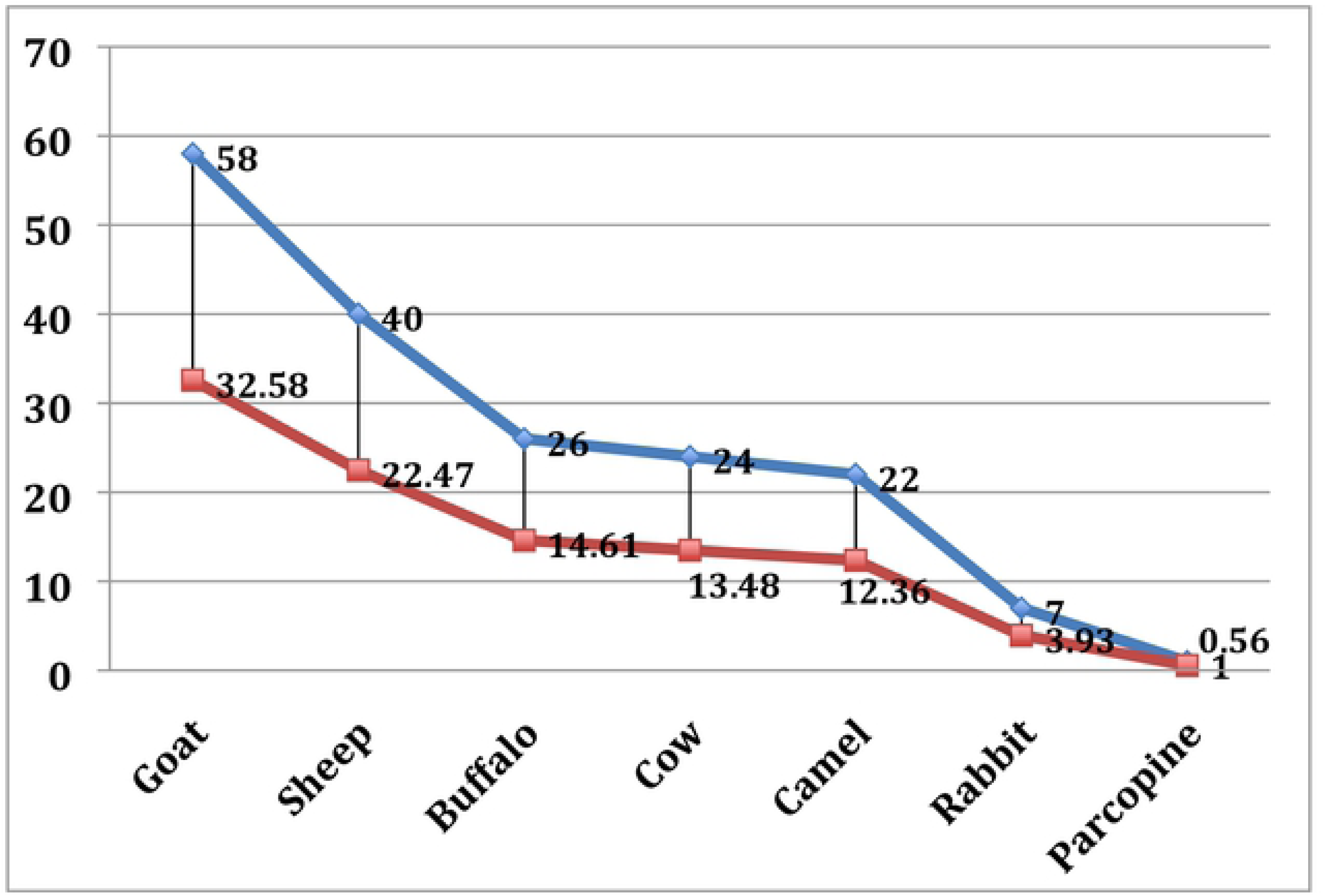
Animals Preferences of grasses in the study area

**Table 6:** Frequency analysis for palatability, parts used for eating and feeding methods and relative abundance of fodder grasses

| Studied parameters | Frequency | Valid percent | Cumulative percent |
| --- | --- | --- | --- |
| Co, Bu, Sh, Go, Ra | 1 | 1.64 | 1.64 |
| Co, Bu, Sh, Go | 6 | 9.84 | 11.48 |
| Co, Bu, Sh, Go, Ra | 4 | 6.56 | 18.03 |
| Go, Sh, Co | 3 | 4.92 | 22.95 |
| Go, Sh | 20 | 32.79 | 55.74 |
| Go, Sh, Co, Cm | 1 | 1.64 | 57.38 |
| Co, Bu, Sh, Go, Cm | 11 | 18.03 | 75.41 |
| Bu, Sh, Go | 1 | 1.64 | 77.05 |
| Co, Bu | 3 | 4.92 | 81.97 |
| Go | 9 | 14.75 | 96.72 |
| Sh | 2 | 3.28 | 100 |
| **Total** | **61** | **100** |  |
| Whole plant | 42 | 68.85 | 68.85 |
| Leaves | 4 | 6.56 | 75.41 |
| Juvenile | 2 | 3.28 | 78.69 |
| Aerial, whole plant at Juvenile | 1 | 1.64 | 80.33 |
| Aerial, Juvenile | 2 | 3.28 | 83.61 |
| Aerial and leaves | 2 | 3.28 | 86.89 |
| Aerial | 8 | 13.11 | 100.00 |
| **Total** | **61** | **100** |  |
| Fo | 31 | 50.82 | 50.82 |
| Fo,For, Mf | 7 | 11.48 | 62.30 |
| Fo, Mf | 21 | 34.43 | 96.72 |
| Fo,Mf | 2 | 3.28 | 100.00 |
| **Total** | **61** | **100** |  |
| Abundant | 30 | 49.18 | 49.18 |
| Common | 13 | 21.31 | 70.49 |
| Frequent | 9 | 14.75 | 85.25 |
| Occasional | 5 | 8.20 | 93.44 |
| Rare | 4 | 6.56 | 100 |
| **Total** | **61** | **100** |  |

Key: Co (Cow), Bu (Buffalo), Sh (Sheep), Go (Goat), Ra (Rabbit), Cm (Camel), Fo (Fodder), For (Forage) , Mf (Mix Fodder)

### Role of the fodder species on milk production

Ten out of the 80 interviewed shepherds (based on the informant knowledge) were randomly sampled to analyze more in detail the role of fodder species on the milk production. We focused our attention on the shepherds because, during the interviews, they showed a deeper knowledge about the species influencing quantity and quality of milk. According to them, *Cynodon dactylon* was the favorite species for the milk production (6.46 SI, 0.6460 CS) followed by *Cymbopogon jwarancusa* (5.133 SI, 0.5133 CS). *Cymbopogon jwarancusa* was also reported to give a peculiar aroma, increasing the milk’s value. *Sorghum* sp. was the third most salient species (5.121 SI, 0.5121 CS) (Table 7). This findings were confirmed when we extended our analysis to all the informants. According to the results of the ANTHROPAC frequency analysis, ranking the plants in the order of their citation frequency (Fig. 9), *Cynodon dactylon* had 73.21% frequency of milk production, following by *Cymbopogon jwarancusa* (70.54%) and *Sorghum* sp. (67.86%).

**Fig. 9:**
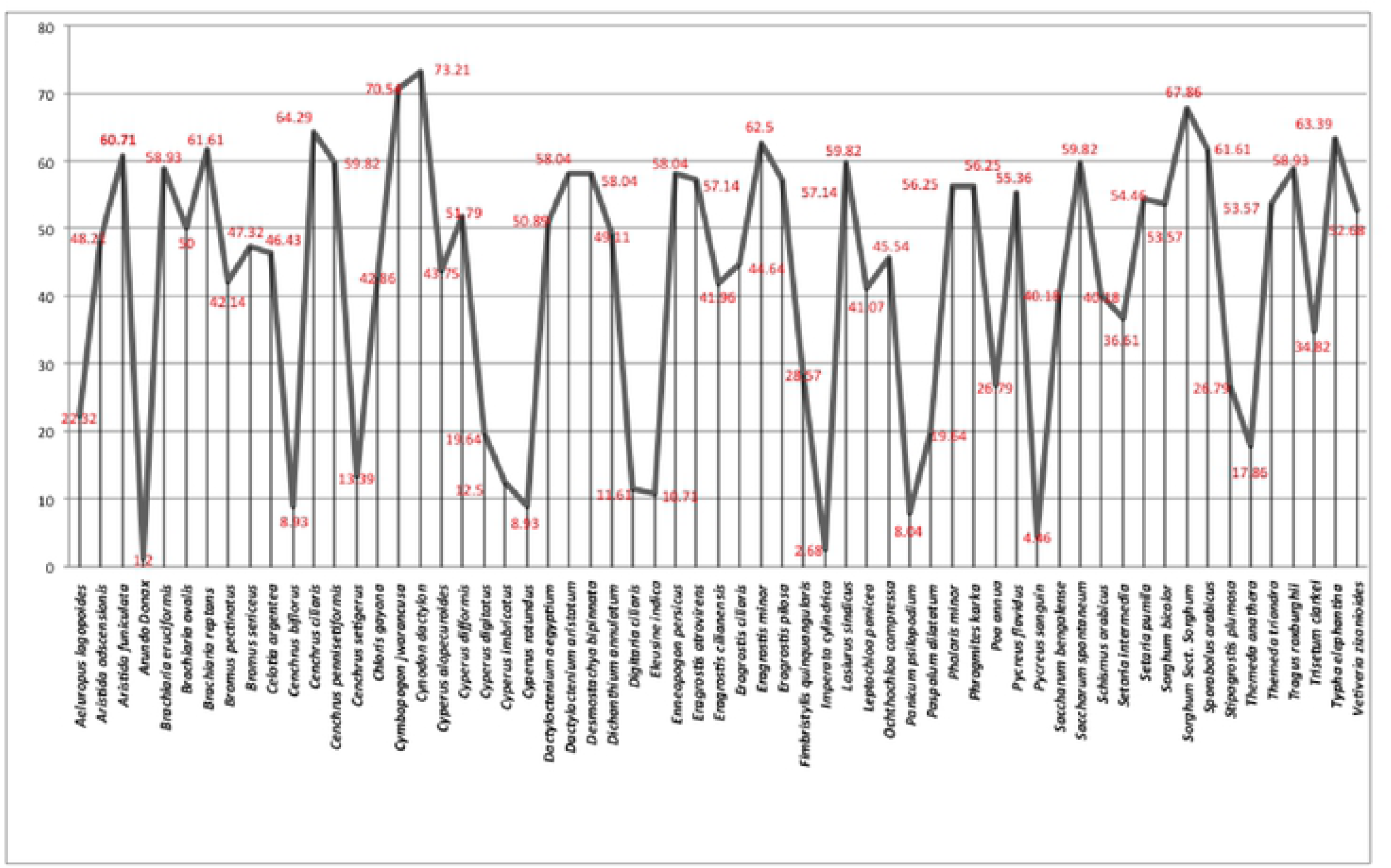
Frequency of milk producing species according to informants ranking

**Table 7:**
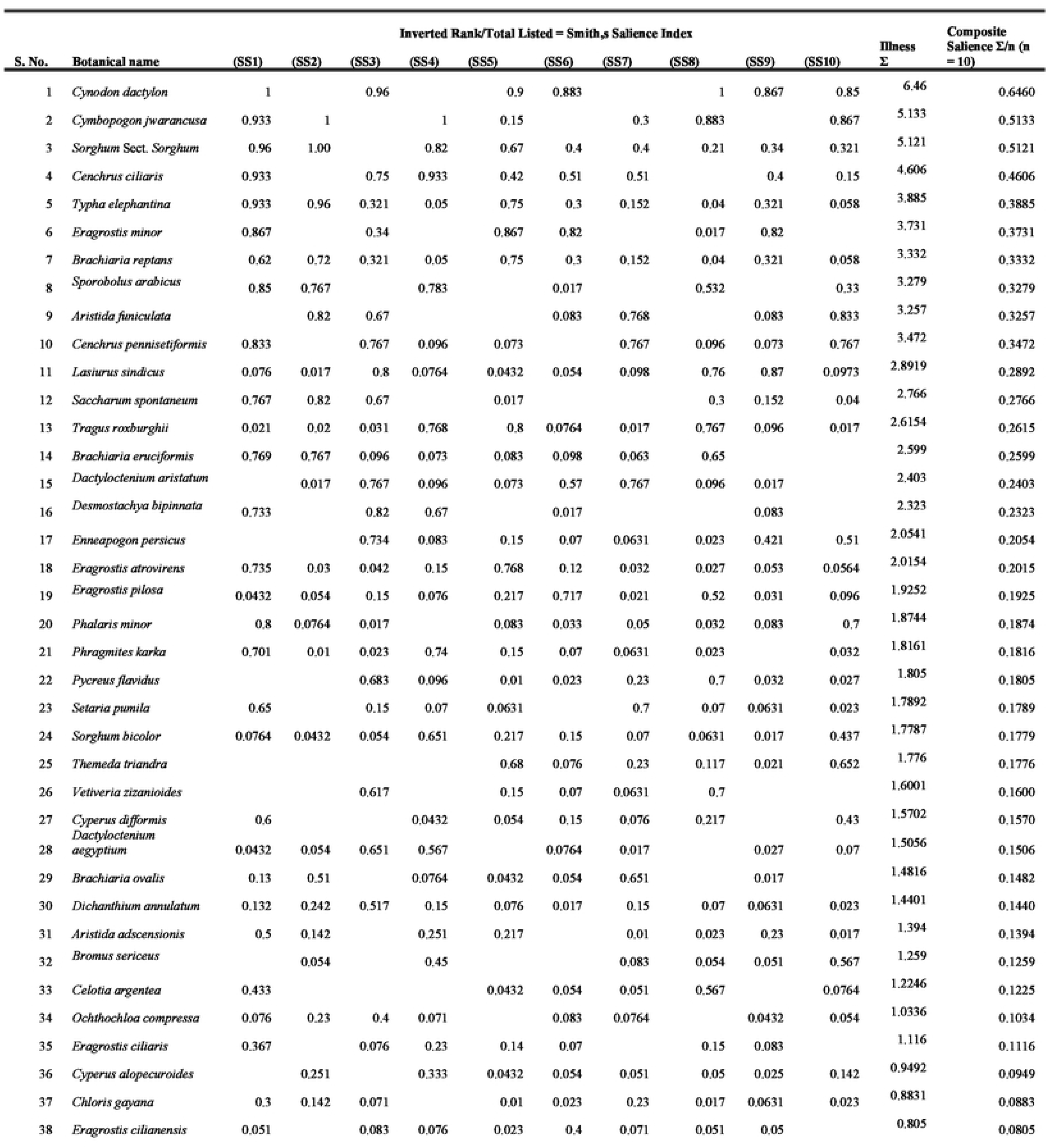
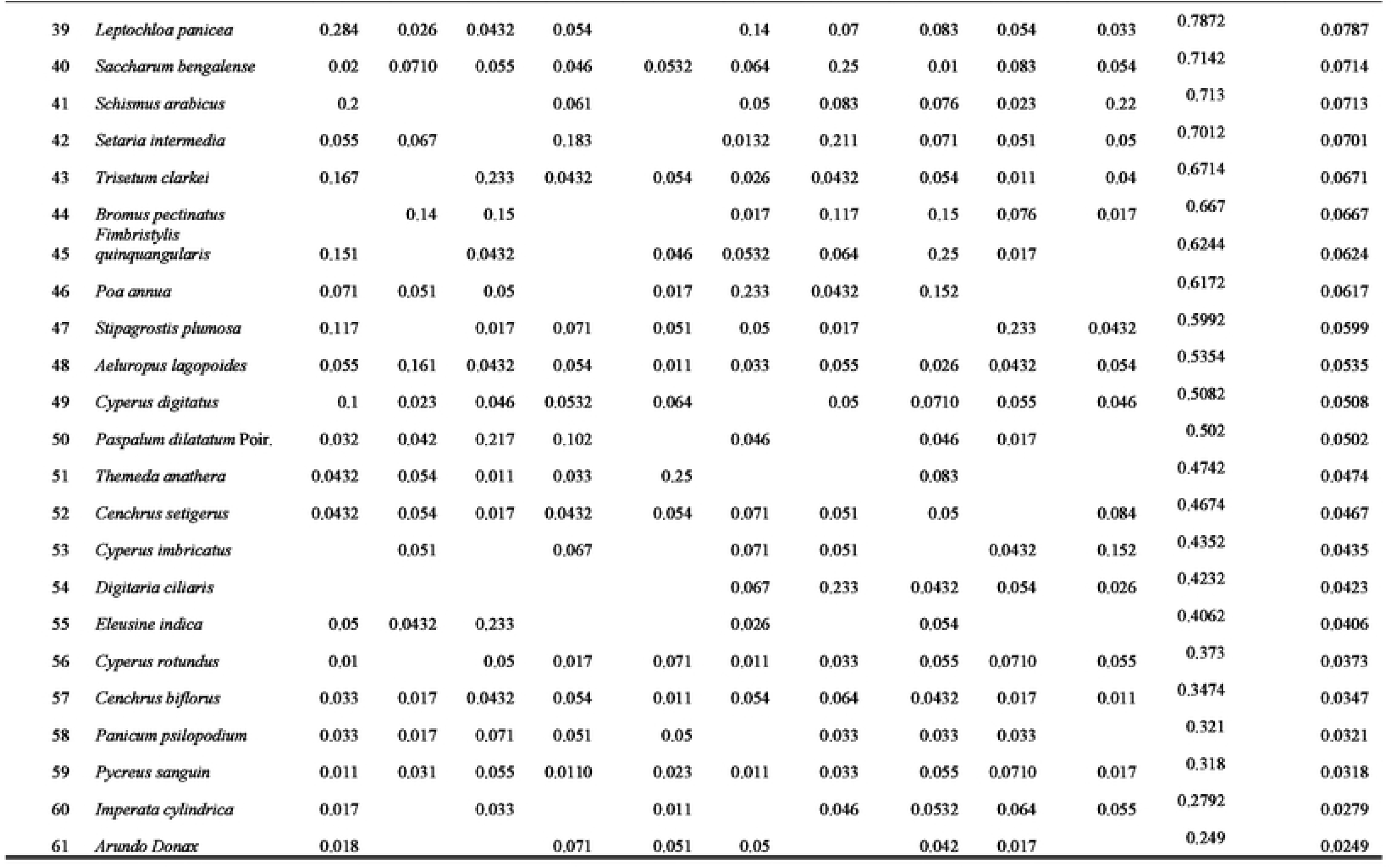
Results of ANTHROPAC analysis of overall salience index of milk producing species

### Relative abundance and seasonal availability

Relative abundance analysis showed that most of the cited species (55%) were abundantly present in the study area and most of them belonged to the priority Group A (Fig.10). 13.39% of the species were available in August and in October while 12.54 % were available in July. In Pakistan, July, August and October are months characterized by monsoon rains fostering the grass biomass development (Fig.11).

**Fig. 10.**
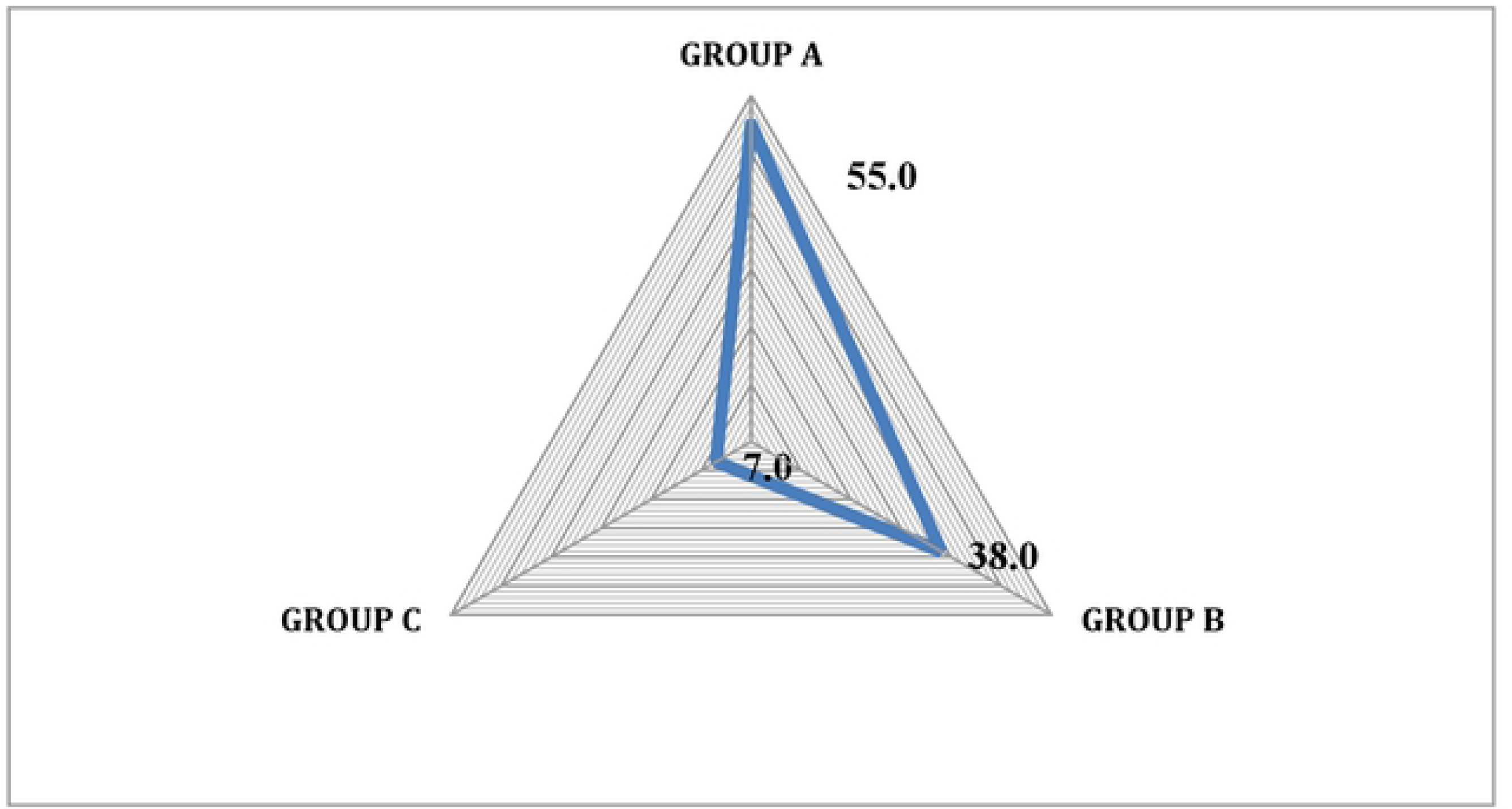
Percentage of species in each group

**Fig. 11:**
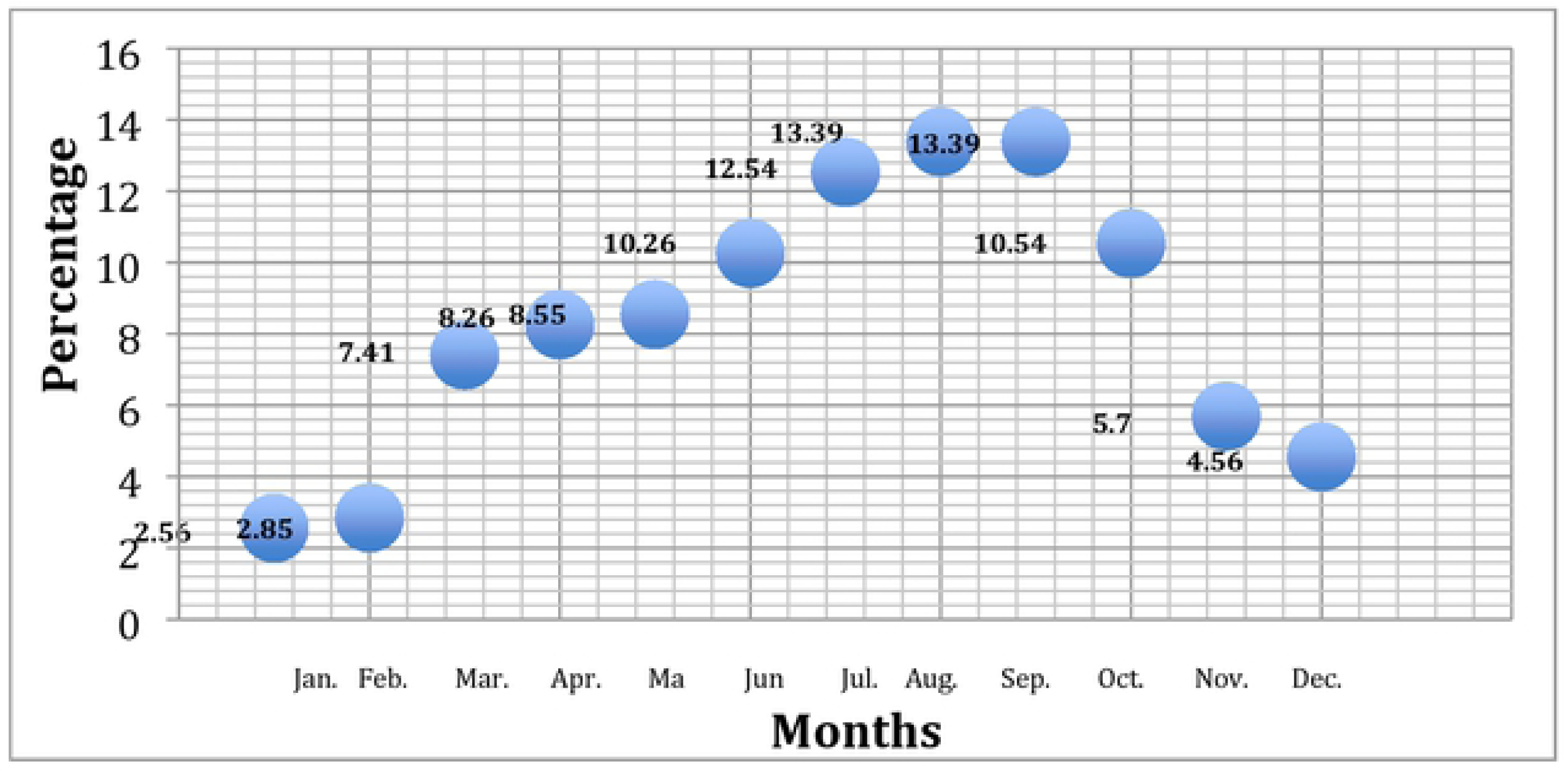
Availability of the grasses in the study area

### People use Livestock for improving their economic life

Livestock production makes the main contribution to agriculture value-added services in the study area. Ten local informants were asked to rank animals from one to five on the basis of their economic value. Milk production is the major income source for people living in the Thal desert; mostly person raised cows and buffaloes more for milk production as compare to raise camels or goats (Fig. 12). Goats, sheep, buffaloes and cows are also raised for meat production. During religious celebrations (such as pilgrimages and *Eid ul Azha*) shepherds and farmers take livestock to the local market for sale and this is another major income source as also shown in [39]. Skin from sheep, buffaloes, cows and camels are also an other way of earning, people sale the animal skin for making leather goods; teeth and bones are used for making different objects (e.g. buttons, jewelry and decoration pieces) (Fig. 12). Dung of buffaloes and cows is dried and used as fuel or, fresh, as a natural fertilizer to improve the soil fertility. Ox, buffaloes and sometimes camels are used for ploughing. Camels are commonly used for transportation in desert areas.

**Fig. 12:**
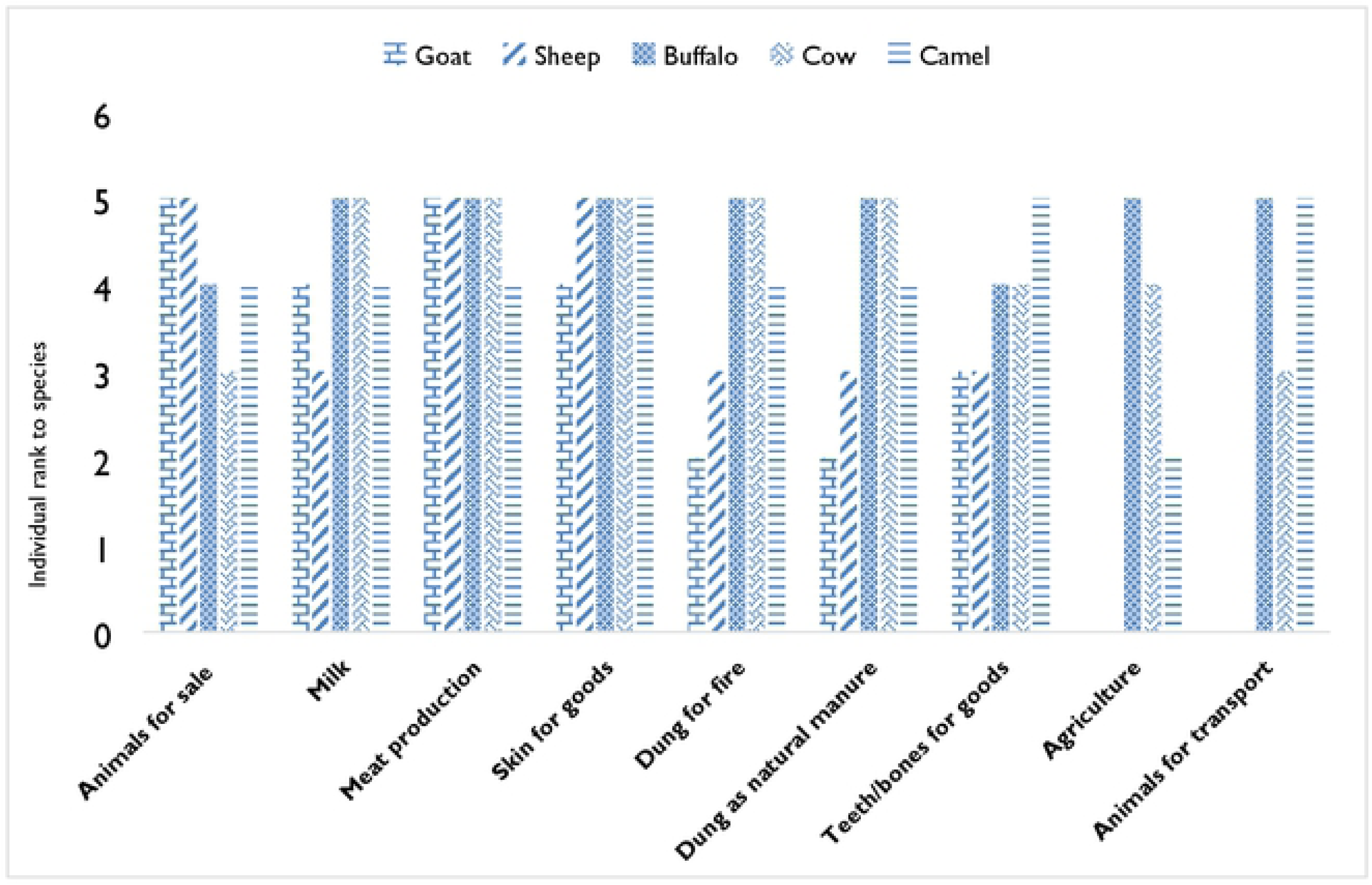
People use Livestock for improving their economic life in Thal Desert

### Indigenous uses and Ethno-veterinary uses of grasses

Eighteen of the 61 reported species were locally used in ethno-veterinary practice. *Cymbopogon jwarancusa* was the most cited veterinary grass (48) and was reported to heal infertility and skin diseases in ruminants (Table 8). Other species (*Cenchrus* spp*., Arundo donax, Desmostachya bipinnata, Dichanthium annulatum, Digitaria ciliaris*, *Eleusine indica, Eragrostis* spp*., Saccharum spontaneum*) were frequently reported to treat urinary and digestive diseases in livestock. As similar results shown in different studies [12, 16, 40]. Urinary and digestive diseases were the most frequently reported disorders; this finding is probably due to the sandy nature of the soil, causing the accumulation of sand-laden feed material in the digestive apparatus and in the urinary tract of livestock.

**Table 8:** Grasses use in Ethnoveterinary and Ethnobotanical

| S. No. | Botanical name | Ethnobotanical Uses | Ethno veterinary uses |
| --- | --- | --- | --- |
| 1 | *Aeluropus lagopoides* (L.)<br>Thwaites | Fuel | --- |
| 2 | *Aristida adscensionis* L | --- | Controls itching |
| 3 | *Arundo Donax* L. | --- | Gastrointestinal |
| 4 | *Arundo Donax* L. | Fencing, inkpot pen, hollow stem for announcement | --- |
| 5 | *Brachiaria ovalis* Stapf | Fuel | --- |
| 6 | *Bromus pectinatus* Thunb. | Fuel | --- |
| 7 | *Bromus sericeus* Drobov | Fuel | --- |
| 8 | *Cenchrus biflorus* Roxb. | --- | Diuretic |
| 9 | *Cenchrus ciliaris* L. | Fuel | Diuretic |
| 10 | *Cenchrus pennisetiformis*<br>Hochst. & Steud. | Fuel | --- |
| 11 | *Cenchrus setigerus* Vahl | Fuel | Diuretic |
| 12 | *Cymbopogon jwarancusa*<br>*subsp. jwarancusa* (Jones)<br>Schult. | Fumigant for measles, matrices (Chatai) for typhoid, root extract for typhus fever and cough, Seeds for chicken pox, roof thatching, roots khass for washing domestic pots/utensils | Fumigant for skin diseases, fragrance in milk, Diuretic and improve fertility in bull |
| 13 | *Cynodon dactylon* (L.)<br>Pers. | Remove pimples, feet burning sensation, fever | Paste of leaves controls dysentery and anti-inflammatory to wounded areas of animal's body |
| 14 | *Cyperus digitatus* Roxb. | Fuel | --- |
| 15 | *Cyperus rotundus* L. | Fuel | Antidiarrheal and gur function stabilizer |
| 16 | *Dactyloctenium aegyptium* (L.) Willd. | --- | Used to reduce after birth abdominal pains |
| 17 | *Desmostachya bipinnata* (L.) Stapf. | Broom making, Fuel | Digestive disorders, Dysentery |
| 18 | *Dichanthium annulatum* (Forssk.) Stapf | Fuel | Digestive disorders |
| 19 | *Digitaria ciliaris* (Retz.)<br>Koel | Fuel |  |
| 20 | *Eleusine indica* (L.)<br>Gaertn. | --- | Cure digestive disorders |
| 21 | *Eragrostis minor* Host | --- | Digestive disorders |
| 22 | *Eragrostis pilosa* (Linn.)<br>P. Beauv. | --- | Help to cure contusion |
| 23 | *Imperata cylindrica* (L.)<br>Racuschel. | --- | Fumigant for Piles |
| 24 | *Leptochloa panicea*<br>(Retz.) Ohwi | Fuel | --- |
| 25 | *Phragmites karka* (Retz.)<br>Trin. ex Steud. | Writing pen (Qalam) trunk, thatching<br>of roof, and fuel source, shoes making | --- |
| 26 | *Pycnus flavidus* (Retz.) T.<br>Koyama | Fuel | --- |
| 27 | *Saccharum bengalense*<br>Retz. | Culms used for making matrices,<br>chairs (Morrhe), hand fan, cages<br>(Pinjra), brooms (Jhaaru), etc. Leaves<br>used for making matrices (Chatai).<br>Leaf sheaths beaten to make strong<br>ropes (Rassi) | Leaves used to treat oral<br>problems of ruminants |
| 28 | *Saccharum spontaneum* L. | Leaves Decoction for stoppage of<br>urination (Micturition), fuel, culm<br>used for making cages, roof thatching<br>(Patalan) and ornamental goods.<br>Leaves woven to make matrices | Root help to relieve in<br>inflammation and urinary<br>problems |
| 29 | *Sorghum bicolor* (Linn.)<br>Moench. | Fuel | Wounds, fever, anemia<br>and constipation |
| 30 | *Sorghum* Sect. *Sorghum*<br>Subsect. *Arundinacea*<br>Moench. | Fuel | --- |
| 31 | *Sporobolus arabicus*<br>Boiss. | Fuel | --- |
| 32 | *Tragus roxburghii*<br>Panigrahi | Fuel | --- |
| 33 | *Typha elephantina* Roxb. | Fuel, roof thatching, ropes, matrices,<br>inflorescence medicinally importance<br>and shoes making | --- |

### Conclusion

The present study is the detailed inventory of 61 indigenous grass species used for fodder and ethno veterinary in Thal district of Southern Punjab Pakistan. The data about grasses was obtained from 232 local informants belonging to different age groups and professions these informants ranked *Cymbopogon jwarancusa* and *Cynodon dactylon* as most preferred grass species. The present study provides an inventory, list of plant parts and diversity in palatability and feeding behavior of these grasses. The data analysis highlighted the possible motives behind the greater acceptability ratio of high priority fodder grasses i.e. diversity in their palatability for major ruminant species, abundant availability in the study area and versatile feeding methods. This data enriched study is not only significant for the conservation of ethnobotanical knowledge but also it may help in facilitating the sustainable livestock feeding for ruminants. Subsequently, the information may play a major role in improving the livelihood of smallholder farmers. Furthermore, it is the first study, which use Smith’s salience index and Composite Salience index to authenticate and validate the collected information. Blend of traditional and scientific knowledge is essentially required to produce worthwhile criterion for selecting these fodder grasses. If some of the grasses show promising nutritional and pharmacological value, then relevant policy marker should take necessary steps to conserve the area and the species. It should not only beneficial for the pharmaceuticssal companies; it will also help to boost up the economy of the country.

## Declaration

### Acknowledgements

We acknowledge all the informants of the study area to for their support and hospitality, We are really very grateful to Director of Camel form Dr. Asharaf and Director of Mani form Dr. Razaq for their support and giving permission to explore the area for data collection. I am very thankful to HEC Pakistan for their kind support for providing travel grant.

## Availability of data and materials

Voucher specimens were submitted to the Herbarium of PMAS-AAUR Pakistan for forthcoming uses (Table 3).

## Authors’ contributions

The ethnobotanical survey and fodder grass sample collection were done by SH, QR, and QFM. QFM did the statistical analysis and SH wrote the manuscript by providing a critical interpretation of the outputs. QR supervised the whole study and helped in identification of specimens. All authors read and approved the final manuscript.

## Ethics approval and consent to participate

### Consent for publication

Not applicable.

